# Fluorescent organelle markers in *Cryptococcus neoformans*: a versatile toolkit for live-cell subcellular localization

**DOI:** 10.64898/2026.03.03.709387

**Authors:** Yeseul Choi, Yong-Sun Bahn, Joseph Heitman

## Abstract

Understanding how intracellular organelles are organized and dynamically remodeled is essential for elucidating fungal growth, differentiation, and pathogenicity. The human fungal pathogen *Cryptococcus neoformans* exhibits remarkable morphological diversity, yet the contribution of intracellular organelles to adaptation across environmental and host-relevant conditions has not been systematically examined. Here, we establish a comprehensive toolkit of fluorescent organelle marker plasmids and strains expressing mCherry or GFP fusions, enabling high-resolution live-cell imaging of ten major subcellular compartments: the nucleolus, endoplasmic reticulum (ER), Golgi apparatus, mitochondria, peroxisome, endosome, autophagosome, vacuole, the plasma membrane, and processing bodies (P-bodies). With this platform, we analyzed organelle organization under host-relevant conditions, including elevated temperature, 5% CO_2_, and capsule- and melanin-inducing media, as well as throughout sexual development from zygote formation to basidiospore production. We further show that the toolkit supports colocalization analyses via diploid formation or dual-labeling strategies. Collectively, our results reveal condition- and stage-specific remodeling of organelle architecture during vegetative growth and mating. This imaging platform provides a robust framework for investigating how subcellular organization supports fungal adaptation, development, and virulence, and for inferring the functions of uncharacterized genes from their spatial localization.

## Introduction

*Cryptococcus neoformans* is an opportunistic fungal pathogen that causes life-threatening meningoencephalitis, particularly in immunocompromised individuals such as HIV/AIDS patients and transplant recipients (Bratton et al. 2012). As a model basidiomycete, *C. neoformans* has a dimorphic life cycle, transitioning from a yeast form to filamentous structures, including hyphae, basidia, and basidiospores, during sexual development (Stempinski et al. 2023). It also produces key virulence factors such as the polysaccharide capsule and polyphenolic melanin (Wang et al. 1995) and is thermotolerant, enabling survival in diverse hosts, including amoebae, macrophages, and microglia (Casadevall et al. 2019; Mansour et al. 2014; Mohamed et al. 2023). Together, these traits reflect a remarkable capacity to adapt to environmental and host-derived stresses—a process that relies on tightly coordinated intracellular responses (Kronstad et al. 2011). The organization and dynamics of subcellular organelles are likely central to these responses, yet they remain poorly characterized in *C. neoformans*, leaving a major gap in our understanding of how intracellular architecture supports fungal virulence and adaptation.

Eukaryotic cells rely on compartmentalization to coordinate essential processes, including metabolism, signaling, stress responses, and molecular turnover. These functions are mediated by both membrane-bound and membrane-less organelles, whose spatial organization within the cytoplasm is tightly regulated rather than random (Zhao and Zhang 2020). Organelle localization is maintained by cytoskeletal networks, motor proteins, and membrane tethers, ensuring efficient trafficking and specially restricted signaling (Hammer and Sellers 2011; Pope and Prade 2025). Even membrane-less assemblies, such as P-bodies and centrosomes, occupy defined cellular locations through microtubule-guided positioning, consistent with their roles in RNA regulation and genome organization (Aizer et al. 2008; Keating et al. 1997). Thus, the spatial positioning and dynamic remodeling of organelles constitute core regulatory features of eukaryotic cell biology.

Recent studies further emphasize that protein function is inherently linked to spatial positioning within the cell, and that dynamic relocalization across compartments can fundamentally alter cellular behavior. A recent review by Sigaeva et al (Sigaeva et al. 2026). highlights subcellular localization as a critical determinant of protein function, emphasizing that spatial positioning within the cell is not merely structural but functionally instructive. Beyond protein distribution, emerging evidence indicates that mRNAs encoding these proteins are themselves spatially regulated, with localization and translational control enabling precise spatiotemporal coordination of protein synthesis. Although the mechanisms governing mRNA localization are not yet fully understood, these observations underscore that spatial regulation operates at multiple levels within the cell. Importantly, proteins are not confined to a single static location but often occupy multiple compartments, and changes in localization, without changes in overall abundance, can substantially influence protein activity and functional output (Christopher et al. 2025; Hein et al. 2025). However, in *C. neoformans*, the spatial organization and remodeling of organelles and associated proteins remain poorly characterized.

Among the subcellular organelles in *C. neoformans*, several compartments play pivotal roles in virulence and adaptation to host-associated stresses by coordinating metabolism, trafficking, and signaling. The ER and Golgi apparatus drive secretion of virulence-associated molecules, including capsule polysaccharides and cell wall-modifying enzymes, through the exocytic pathway (Mota et al. 2025). Mitochondria and peroxisomes support energy production and oxidative stress responses (Kretschmer et al. 2012), while vacuoles function in ion homeostasis, autophagy, and nutrient storage (Li and Kane 2009). Endosomes and autophagosomes contribute to nutrient and ion uptake, capsule elaboration, and virulence through ESCRT- and ATG-dependent pathways (Hu et al. 2015; Ding et al. 2018). The plasma membrane mediates environmental sensing and extracellular vesicle-dependent export of virulence factors (Xiao et al. 2025). The nucleolus regulates ribosomal biogenesis and has been linked to stress-responsive cell cycle control, including the G2 arrest associated with titan cell formation, which promotes adaptation to host environments (Altamirano et al. 2021). P-bodies are induced in *C. neoformans* under host-like heat stress, consistent with roles in stress-responsive mRNA processing (Kozubowski et al. 2011). Despite clear evidence that cellular organelles contribute to virulence and stress adaptation, how organelle architecture and dynamics are remodeled in response to host- and environment-derived cues remain poorly defined.

In this study, we developed and employed a genetically encoded organelle marker system to perform high-resolution live-cell imaging of *C. neoformans* across a range of physiological and host-mimicking conditions, including elevated temperature with or without 5% CO_2_, capsule- and melanin-inducing media, co-culture with human brain endothelial cells, and mating-inducing media to analyze the sexual cycle. This approach enabled us to define organelle-specific responses to diverse stimuli, underscoring the importance of subcellular compartmentalization in fungal adaptation and virulence. The system also supports colocalization studies through diploid formation or dual-labeling in haploid strains, facilitating functional analysis of candidate proteins within defined spatial contexts. Together, this platform provides a robust framework to investigate organelle dynamics and to infer functions of uncharacterized genes based on subcellular localization.

## Materials and methods

### Strains, media, and growth conditions

All *Cryptococcus* strains utilized in this study are listed in Table S1. Unless otherwise noted, cells were cultured in yeast extract (1%)–peptone (2%)–dextrose (2%) (YPD) medium at 30°C with shaking at 200 rpm. YPD was supplemented with nourseothricin (100 μg/ml), G418 (50 μg/ml), or hygromycin B (150 μg/ml) as needed to select *C. neoformans* transformants generated by biolistic particle delivery or CRISPR-based methodology. For stress or host-relevant conditions, cultures were incubated at 37°C, with 5% CO_2_ as indicated. Capsule-inducing conditions were achieved in DMEM, and melanin production was induced on agar containing L-DOPA. For mating and hyphal development, cells were spotted onto V8 agar (pH=5.0) or MS medium and incubated in the dark at room temperature for the indicated times.

### Construction of a versatile fluorescent tagging plasmid

For construction of pHYG_SH_P_H3__mCherry, the safe haven (SH) flanking sequence and the constitutive histone *H3* (P*_H3_*) promoter were PCR-amplified. To introduce linearization restriction enzyme sites (AscI, PacI, and BaeI) within the SH region, these sites were incorporated by double-joint PCR during fragment assembly. The amplified SH fragment and *H3* promoter fragment were cloned into pHYG_mCherry at the NotI site by Gibson assembly (New England Biolabs, E2611). The NotI site was retained immediately upstream of the mCherry–linker sequence to facilitate subsequent insertion of target genes. After the backbone vector was generated, marker genes or genes of interest were cloned into the retained NotI site by Gibson assembly to express proteins as C-terminal mCherry fusions under control of the *H3* promoter.

For construction of pHYG_SH_P*_H3_*_GFP, the SH fragment, *H3* promoter, GFP coding sequence, and the *HOG1* terminator were fused together by double-joint PCR using Phusion High-Fidelity DNA Polymerase Master Mix (Thermo Fisher Scientific, F-548). The fused fragment was inserted into the pHYG backbone plasmid linearized by NotI digestion using NEBuilder HiFi DNA Assembly Master Mix (New England Biolabs, E2621). As in the mCherry construct, a single NotI site was retained immediately upstream of the GFP coding sequence to enable insertion of target genes for fluorescent tagging.

To generate a selectable marker variant, the hygromycin resistance cassette (*HYG*) in pHYG_SH_P_H3__GFP was replaced with a neomycin resistance cassette (*NEO*). The plasmid was digested with BstXI to remove the *HYG* marker, and the *NEO* cassette was PCR-amplified and cloned into the digested vector using NEBuilder HiFi DNA Assembly Master Mix. Genes of interest were subsequently inserted into the retained NotI site of pHYG_SH_P_H3__GFP or pNEO_SH_P_H3__GFP by NEBuilder HiFi DNA Assembly Master Mix.

### Construction of mutant strains

For generation of mCherry marker strains, plasmids carrying pHYG_SH_P_H3__Gene_mCherry were linearized with PacI and introduced into the H99α strain by biolistic transformation (Kim et al. 2009). To generate GFP marker strains, pHYG_SH_P_H3__Gene_GFP plasmids were similarly linearized with PacI and introduced into the KN99**a** strain using transient CRISPR–Cas9–coupled electroporation (TRACE) (Lin et al. 2015; Fan and Lin 2018). Correct targeted integration of each marker construct at the SH locus was verified by diagnostic PCR using primer pairs annealing upstream and downstream of the integration site. Primer sequences used for diagnostic PCR are listed in Table S2.

### Quantitative reverse transcription-PCR (qRT-PCR) analysis of gene expression

Wild-type and mutant strains were cultured overnight in YPD medium at 30°C and then diluted to an OD_600_ of 0.2 in fresh medium. Cultures were grown to mid-log phase (OD_600_ = 0.8) and harvested by centrifugation. Cell pellets were frozen and lyophilized. Total RNA was isolated using the mirVana™ miRNA Isolation Kit (Thermo Fisher Scientific, AM1561) according to the manufacturer’s instructions. For cDNA synthesis, total RNA was adjusted to 2 µg with DNase- and RNase-free water, mixed with anchored oligo(dT)_23_ primers (Sigma-Aldrich, O4387), and incubated at 65°C for 5 min. The reaction was supplemented with RNase inhibitor (Applied Biosystems, N8080119), dNTPs, and reverse H minus reverse transcriptase (Thermo Fisher Scientific, EP0752), followed by incubation at 50°C for 60 min. Reverse transcription were terminated by heat inactivation at 85°C for 10 min. Quantitative RT-PCR was performed with gene-specific primers, with *ACT1* serving as an internal reference gene. Primer sequences are listed in Table S2.

### Diploid strain construction

To generate diploid strains, *MAT*α and *MAT***a** cells were cultured overnight in YPD medium at 30°C. Cultures were washed twice with phosphate-buffered saline (PBS) and adjusted to 1x10^8^ cells/ml. Equal volumes of *MAT*α and *MAT***a** cells suspensions were mixed and spotted onto V8 agar (pH=5). Plates were incubated in the dark at 25°C for 24 h. The mating spot was then scraped, resuspended in 1 ml of distilled water, and plated onto selection media (supplemented with G418 and hygromycin B). Colonies recovered on selection plates were streaked onto fresh selective media to confirm stable diploid growth.

### Visualization of subcellular localization of mCherry- or GFP-tagged cellular marker proteins

mCherry- and GFP-tagged *Cryptococcus* strains were grown overnight in YPD medium at 30°C with shaking. Cultures were then diluted into fresh YPD and grown to mid-log phase (OD_600_=0.8). Cells were harvested by centrifugation and processed immediately for fluorescence microscopy unless otherwise specified. For experiments performed under specific growth or stress conditions, cells were cultured according to the conditions described for each experiment. Fluorescence images were acquired using a Zeiss Axioskop 2 Plus epifluorescence microscope equipped with an Axiocam 305 mono camera. For confocal imaging, cells were mounted on glass slides overlaid with 2% agarose and visualized using an Andor Dragonfly spinning disk confocal microscope.

### Organelle staining

For nuclear staining, cells were fixed in 4% paraformaldehyde containing 3.4% sucrose for 15 min at room temperature. Fixed cells were washed with buffer containing 0.1 M potassium phosphate (KPO_4_) and 1.2 M sorbitol and stained with 10 µg/mL Hoechst 33342 (Thermo Fisher Scientific, H1399) for 30 min in the dark. Cells were then washed and imaged by fluorescence microscopy.

For ER staining, harvested cells were washed twice with Hank’s Balanced Salt Solution (HBSS) (Gibco, 14025092). Cells were fixed as described above and resuspended in HBSS buffer. ER-Tracker dye (Invitrogen, E34251) was added to a final concentration of 1 µM, and cells were incubated at 30°C for 30 min in the dark. After incubation, cells were washed twice with HBSS prior to imaging.

For mitochondrial staining, harvested cells were incubated with 50 nM Mito-Tracker dye (Invitrogen, M7514) for 15 min at room temperature in the dark. Cells were washed with PBS and immediately examined by fluorescence microscopy.

### P-body and autophagosome induction

To induce P-body formation, cultures grown to OD_600_ = 0.8 were shifted to 37°C and incubated for 15 min. Cells were then immediately imaged by fluorescence microscopy without fixation. To visualize autophagosome localization under autophagy-inducing conditions, cells were inoculated at an OD_600_ of 0.2 into yeast nitrogen base (YNB) medium lacking amino acids and 2% glucose, supplemented with 1 mM phenylmethylsulfonyl fluoride (PMSF) and 10 µg/mL nocodazole. Cultures were incubated at 30°C for 4 h, harvested, and examined by fluorescence microscopy.

### Vacuole morphology analysis

To induce changes in vacuolar morphology, cells were inoculated at an initial OD_600_ of 0.2 into three different media conditions: 1/5-diluted YPD (hypotonic), standard YPD (isotonic), and YPD supplemented with 1 M NaCl (hypertonic). Cultures were incubated at 30°C and harvested at OD_600_ = 0.8 for fluorescence microscopy.

### Titan cell induction

For titan cell formation, 5 x 10^3^ yeast cells were inoculated in 1 mL of serum-free RPMI 1640 medium (Sigma-Aldrich, R1383) in a 24-well tissue culture plate. Cells were incubated at 37°C with 5% CO_2_ for 24 h without shaking.

### *In vitro* phenotypic analysis

To examine growth characteristics and chemical susceptibility, each strain was cultured in liquid YPD at 30°C for 16 h with shaking. Cultures were serially diluted 10-fold (10⁰ to 10^-4^) and spotted onto YPD agar containing the indicated chemical stressors. Plates were incubated at 30°C for up to 5 days and monitored daily. Representative images were captured to compare relative growth.

To compare protein localization and fluorescence intensity across environmental conditions, 2 mL of overnight-cultures were spotted onto YPD agar and incubated at 30°C, 37°C, or 37°C with 5% CO_2_ for 2 days. Cells were then harvested and imaged by fluorescence microscopy using identical exposure settings. Fluorescence intensity was quantified in Fiji/ImageJ. Cell boundaries were defined using DIC images and applied to the corresponding fluorescence channels. Mean fluorescence intensity per cell was measured after background subtraction. At least 50 cells were analyzed per condition. Values were normalized to the 30°C condition and are presented as mean ± standard deviation (SD).

For melanin assays, overnight-cultures were washed twice with PBS and spotted onto Niger seed agar containing 0.1% glucose. Plates were incubated at 37°C and imaged daily for 1–3 days. For fluorescence analysis, cells were scraped from plates on day 2, resuspended in PBS, and imaged by fluorescence microscopy.

For capsule induction, overnight cultures were washed with PBS and spotted onto RPMI agar, then incubated at 37°C. After 2 days, cells were stained with India ink and examined by differential interference contrast (DIC) microscopy. Capsule thickness was calculated for at least 50 randomly selected cells per strain using the formula: capsule thickness = total cell diameter − cell body diameter. Statistical significance was assessed by one-way ANOVA with Bonferroni’s multiple-comparison test.

### Fluorescence analysis during mating

For fluorescence analysis during mating, *MAT*α and *MAT***a** strains expressing mCherry-or GFP-tagged proteins were mixed at equal cell densities and spotted onto MS medium (Sigma-Aldrich, M5524). Plates were incubated in the dark at room temperature for the indicated times to allow filamentation and sexual development. To visualize zygotes, mating spots were scraped after 3 days, resuspended in distilled water, and examined by fluorescence microscopy. Hyphae and basidial structures were imaged directly from excised agar blocks.

## Results

### Design and validation of a standardized fluorescent organelle marker platform in *C. neoformans*

To establish a robust, standardized platform for visualizing subcellular organelles in *C. neoformans*, we constructed a series of plasmids designed for targeted integration at the genomic safe haven locus (Arras et al. 2015). Each plasmid was designed to enable NotI-mediated insertion of a gene of interest and to drive expression from the constitutive histone *H3* promoter (Fig. 1a and b). Inserted genes were fused at their C termini to either mCherry or GFP and terminated with the *HOG1* transcriptional terminator, with hygromycin resistance as the selectable marker (Fig. 1a).

**Fig. 1.**
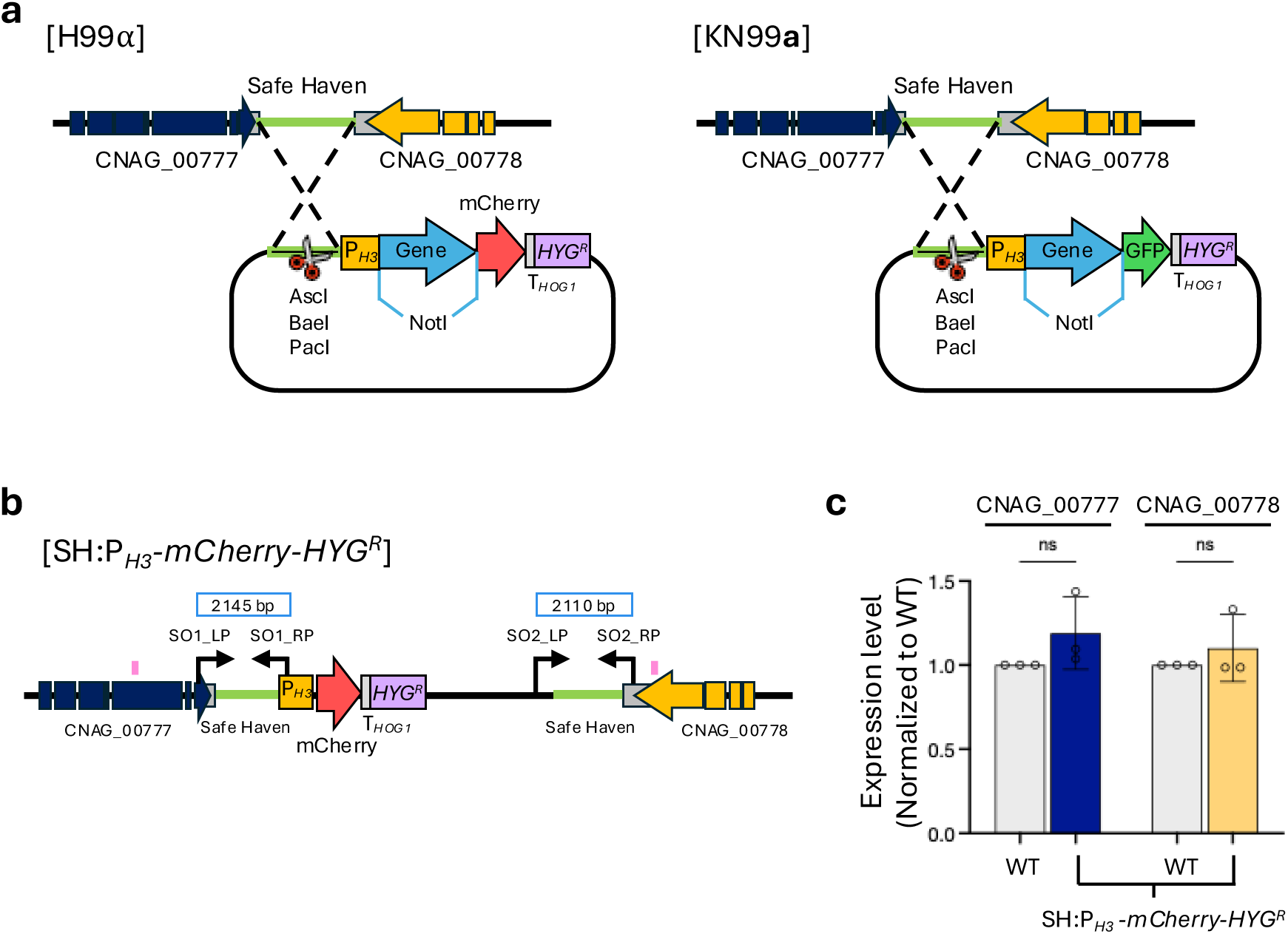
Strategy for construction and validation of fluorescent organelle marker strains in *C. neoformans*. a) Schematic representation of the strategy used to generate fluorescent organelle marker strains. Target genes were fused to GFP or mCherry under the control of the constitutive *H3* promoter and integrated into the safe haven locus by homologous recombination. mCherry-tagged constructs were integrated into the H99α background, and GFP-tagged constructs into the KN99**a** background. b) Diagrams illustrating the insertion of the P*_H3_-gene-mCherry-HYG* or P*_H3_-gene-GFP-HYG* cassette into the safe haven locus between CNAG_00777 and CNAG_00778 via homologous recombination. Correct targeted integration was confirmed by diagnostic PCR across both recombination junctions: the upstream junction was verified using primer pairs SO1_LP and SO1_RP, and the downstream junction using SO2_LP and SO2_RP. The regions highlighted in pink indicate the qRT-PCR regions within each gene. c) Gene expression analysis. Expression levels of CNAG_00777 and CNAG_00778 were measured by qRT-PCR. RNA was extracted from cells harvested at the OD_600nm_ = 0.8. Statistical analysis was performed using one-way ANOVA. Error bars represent SD.

Consistent with the original safe haven design (Arras et al. 2015), the plasmid backbone contains AscI, BaeI, and PacI restriction sites to enable linearization prior to transformation. Targeted integration was confirmed by diagnostic PCR with primers flanking the upstream (CNAG_00777) and downstream (CNAG_00778) regions of the safe haven locus (Fig. 1b, Supplementary Fig. 1a). Importantly, qRT-PCR analysis showed that transcript levels of both flanking genes were unchanged after plasmid integration, confirming that insertion at the safe haven locus does not perturb expression of neighboring genes (Fig. 1b and c).

### Distinct spatial organization of subcellular organelles in *C. neoformans*

To generate a comprehensive set of organelle markers, we selected 11 genes representing 10 major subcellular compartments, including the nucleolus, P-bodies, mitochondria, ER, Golgi apparatus, vacuole, peroxisome, endosome, autophagosome, and plasma membrane (Fig. 2a and b). Marker selection was guided by three criteria: (i) documented organelle-specific functions in *C. neoformans*, (ii) prior use as validated markers, and (iii) conserved localization of orthologs in the model yeast *Saccharomyces cerevisiae* and other model fungi. For dynamic organelles, including the ER, Golgi apparatus, and endosomes, we selected proteins with non-overlapping localization patterns to ensure unambiguous spatial assignment.

**Fig. 2.**
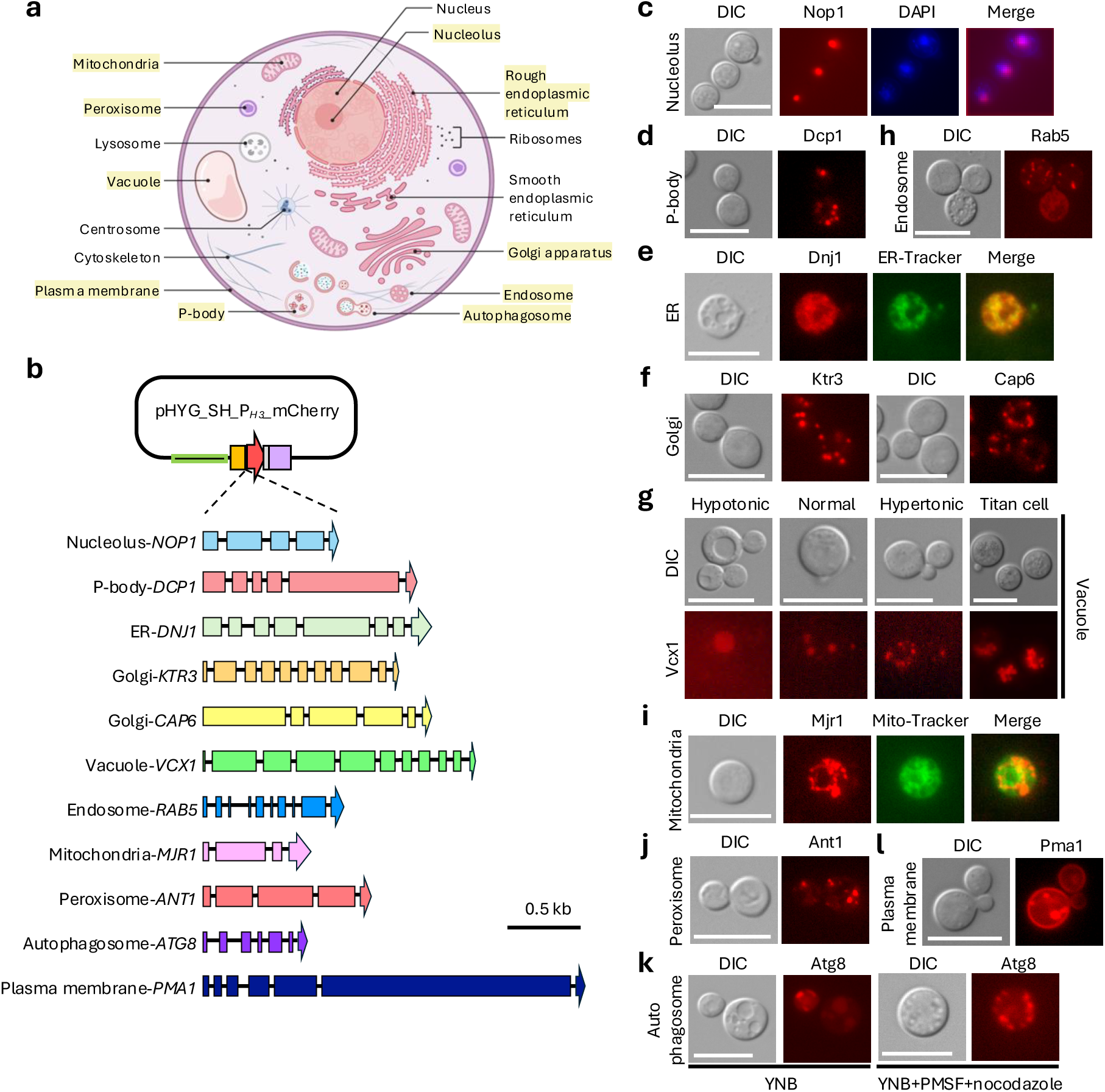
Comprehensive fluorescent organelle marker toolkit for *C. neoformans*. a) Schematic illustration of a *C. neoformans* cell highlighting the major intracellular organelles targeted in this study. The image was created with Biorender. b) Ten representative genes corresponding to distinct intracellular organelles. Each gene was subcloned into the pHYG_SH_P_H3__mCherry vector. Representative organelle marker genes used in this study: *NOP1* (nucleolus), *DCP1* (P-body), *DNJ1* (ER), *KTR3* and *CAP6* (Golgi), *VCX1* (vacuole), *RAB5* (endosome), *MJR1* (mitochondria), *ANT1* (peroxisome), *ATG8* (autophagosome), and *PMA1* (plasma membrane). c-l) Representative fluorescence microscopy images showing the subcellular localization pattern of selected organelle marker in *C. neoformans*. c) Nucleolus marker. Nucleolar localization was visualized using Nop1–mCherry (SH:P*_H3_*-*NOP1-mCherry*, YSB11829). Cells were fixed and stained with Hoechst dye for nuclear visualization. d) P-body marker. To induce P-body formation, cultures harboring Dcp1–mCherry (SH:P*_H3_*-*DCP1-mCherry*, YSB11825) were grown at 30°C to mid-log phase (OD_600nm_ = 0.8), briefly shifted to 37°C for 15 min, and immediately imaged. e) ER marker. The ER network was examined using Dnj1–mCherry protein (SH:P*_H3_*-*DNJ1-mCherry*, YSB11832) following fixation and staining with ER-Tracker dye. f, h, j, and l) Golgi, endosomal, peroxisomal, and plasma membrane markers. Cells featuring mCherry-tagged cellular markers−SH:P*_H3_*-*KTR3-mCherry* (YSC8), SH:P*_H3_*-*CAP6-mCherry* (YSC11), SH:P*_H3_*-*RAB5-mCherry* (YSC9), SH:P*_H3_*-*ANT1-mCherry* (YSB11817), SH:P*_H3_*-*ATG8-mCherry* (YSB11835), and SH:P*_H3_*-*PMA1-mCherry* (YSC4) −were grown at 30°C to mid-log phase (OD_600nm_ = 0.8) and imaged without fixation. g) Vacuole marker. Vacuolar morphology and size variability were assessed using the cells expressing Vcx1–mCherry (SH:P*_H3_*-*VCX1-mCherry*, YSB11822) under multiple growth conditions. Cells were cultured in standard YPD, 1/5-diluted YPD, or YPD supplemented with 1 M NaCl, and vacuoles were additionally examined in titan cell–induced populations. i) Mitochondrial marker. Mitochondrial localization was confirmed by imaging the cells featuring mCherry tagged Mjr1 (SH:P*_H3_*-*DNJ1-mCherry*, YSB11832) following staining with Mito-Tracker dye. k) Autophagosomal marker. Autophagosome formation was monitored using Atg8–mCherry (SH:P*_H3_*-*ATG8-mCherry*, YSB11835) in cells cultured in YNB medium supplemented with PMSF and nocodazole. Scale bar; 10 μm.

For each marker, the full-length coding sequence (including all annotated exons) was cloned into the standardized plasmid backbone (Fig. 2b), enabling C-terminal fluorescent tagging while preserving the endogenous coding sequence. To generate organelle marker strains, constructs were integrated into the *MAT*α H99 strain. Correct integration at the safe haven locus was confirmed by diagnostic PCR with primers spanning the 5’ and 3’ junction flanking regions (Fig. 1b and Supplementary Fig. 1a). For representative mCherry-tagged strains, we further confirmed that marker genes were overexpressed as intended and that transcript levels of genes flanking the safe haven locus were unchanged after integration (Supplementary Fig. 1b). In addition, integration of the organelle markers did not alter stress responses, suggesting that marker integration does not disrupt stress-related phenotypes (Supplementary Fig. 1c).

To validate markers associated with nuclear and cytoplasmic RNA granules, we examined nucleolar and P-body localization under conditions that robustly reveal their defining structural features. Localization of the nucleolar marker Nop1 (Lee and Heitman 2012) was validated by DAPI staining, which showed colocalization of the fluorescent signal with the nucleolar region (Fig. 2c). For P-body markers, we tagged Dcp1, a core P-body component (Kozubowski et al. 2011). Cells grown at 30°C were shifted to 37°C for 15 min prior to imaging, a condition that robustly induced punctate P-body structures (Fig. 2d).

To characterize markers of the secretory and endomembrane system, we analyzed the spatial organization and compartment-specific distribution of ER, Golgi, endosomal, and vacuolar markers. For the ER, we tagged Dnj1, an ER-resident protein (Horianopoulos et al. 2021), and performed colocalization analysis with a green ER-Tracker dye. ER-mCherry overlapped strongly with ER-Tracker staining and were prominently enriched in perinuclear regions (Fig. 2e). For the Golgi apparatus, we independently tagged two genes implicated in Golgi function in *C. neoformans*, *CAP6* and *KTR3* (Thak et al. 2022); both showed clear punctate cytoplasmic localization (Fig. 2f). We evaluated the vacuolar marker Vcx1 (Kmetzsch et al. 2010) under hypotonic and hypertonic conditions, as well as during titan cell induction. As expected, hypotonic growth produced enlarged vacuoles, whereas hypertonic conditions led to fragmented punctate structures (Fig. 2g). In titan cells, multiple vacuoles were observed within individual cells (Fig. 2g). We next examined Rab5, a regulator of early endosomal trafficking (Aboobakar et al. 2011), which displayed punctate cytoplasmic localization consistent with early endosomes (Fig. 2h).

To assess markers of metabolic organelles, we analyzed mitochondrial and peroxisomal localization patterns. For mitochondria, we tagged Mjr1 (Horianopoulos et al. 2020) and performed colocalization analysis with a green Mito-Tracker dye. Mjr1–mCherry largely overlapped with Mito-Tracker staining, while also labeling a more extensive mitochondrial network that extended beyond the Mito-Tracker positive regions (Fig. 2i). For peroxisomes, we tagged Ant1 based on its established role in peroxisomal metabolism (El Magraoui et al. 2014). Ant1 showed characteristic punctate cytoplasmic localization, with multiple discrete foci per cell, consistent with peroxisomal organization (Fig. 2j).

As an autophagosome marker, we tagged Atg8, which localizes to autophagosomal membranes during autophagosome biogenesis in *C. neoformans* (Zhao et al. 2019). To enhance visualization of Atg8-labeled structures, cells were incubated for 4 h in minimal medium containing PMSF and nocodazole, a condition previously shown to promote autophagosome accumulation. Under these conditions, Atg8-positive puncta were readily detected and were frequently enriched near the cell periphery (Fig. 2k). To label the plasma membrane, we tagged Pma1, a plasma membrane H^+^-ATPase that undergoes vacuole-mediated trafficking before reaching the cell surface (Estrada et al. 2015), enabling visualization of both the plasma membrane and vacuolar compartments (Fig. 2l).

Together, these results validate all 11 marker genes representing 10 distinct subcellular compartments as functional, organelle specific, and suitable for visualizing organelle organization in *C. neoformans*.

### Organelle-specific remodeling under host-relevant temperature and CO_2_ conditions

Host-relevant cues such as elevated temperature and CO_2_ are key determinants of adaptation and virulence in *C. neoformans* (Chadwick et al. 2025). Although these conditions trigger broad transcriptional and physiological changes, how individual organelles quantitatively respond to temperature and CO_2_ remains poorly defined. To address this, we systematically quantified organelle-associated fluorescence in cells grown at 30°C and host-like conditions (37°C and 37°C with 5% CO_2_) utilizing our panel of fluorescent organelle marker strains. Fluorescence microscopy was performed under identical acquisition settings across all conditions to enable direct comparison of signal intensity between treatments.

Quantitative analysis revealed that organelle-associated fluorescence responses to elevated temperature and CO_2_ fell into three distinct patterns: organelles responsive to both temperature and CO_2_, organelles selectively responsive to either high temperature or CO_2_, and organelles largely unresponsive to either cue. In the temperature- and CO_2_-responsive group, the nucleolar marker Nop1, the ER marker Dnj1, and the endosomal marker Rab5 exhibited significantly increased fluorescence at 37°C compared with 30°C, with a further increase at 37°C plus 5% CO_2_ (Fig. 3a). These results suggest that nucleolus, ER, and endosomal compartments are particularly sensitive to combined host-like cues of temperature and CO_2_.

**Fig. 3.**
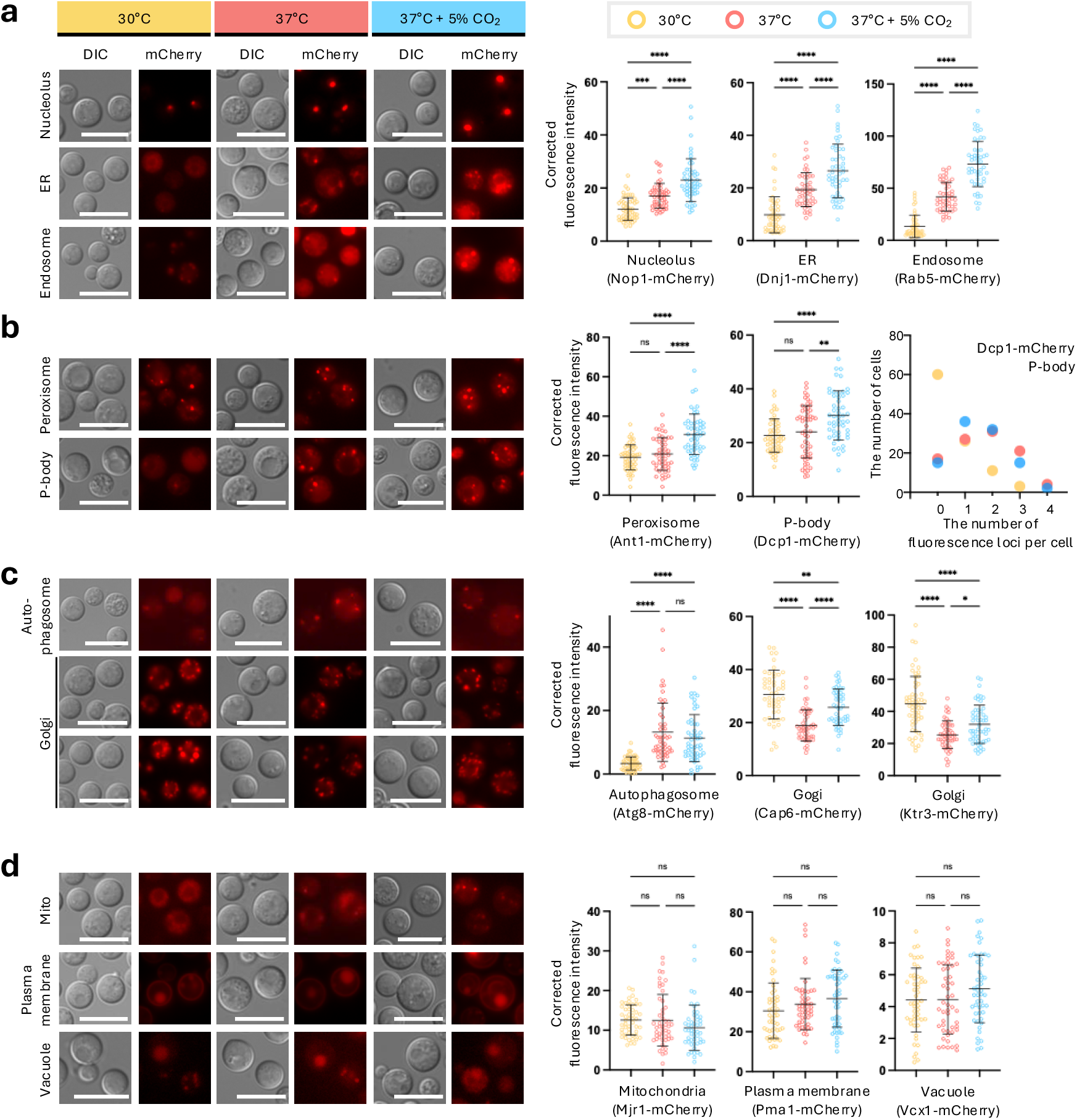
Increased fluorescence intensity of multiple organelle markers under elevated temperature and CO_2_ conditions. a-d) mCherry-tagged organelle marker strains in *C. neoformans* were cultured at 30°C, 37°C, 37°C with 5% CO_2_ and analyzed by fluorescence microscopy. For each organelle marker, cells were imaged under identical acquisition settings to enable direct comparison of fluorescence intensity across conditions. Representative images are shown in the left panels, while the corresponding quantification of fluorescence intensity is shown in the right panels. Fluorescence intensity was measured from 50 individual cells per condition. Statistical significance was determined by one-way ANOVA (*, *p* < 0.05; **, *p* < 0.01; ****, *p* < 0.0001; ns, not significant). Scale bars, 10 µm. b, rightmost panel) Quantification of cytoplasmic Dcp1–mCherry P-body foci per cell in the cells expressing Dcp1–mCherry (*n* = 50 cells per condition).

In contrast, the peroxisomal marker Ant1 and the P-body marker Dcp1 showed little change in fluorescence intensity at 37°C alone but increased markedly under 37°C plus 5% CO_2_ (Fig. 3b), indicating selective responsiveness to elevated CO_2_. Notably, Dcp1 fluorescence was broadly distributed throughout the cytoplasm at 30°C, whereas incubation at 37°C promoted the formation of discrete punctate P-body structures. Under 37°C plus 5% CO_2_, fluorescence intensity increased while puncta were maintained, suggesting that CO_2_ promotes both increased abundance and sustained assembly of P-bodies.

The autophagosome marker Atg8 showed a selective response to elevated temperature. Under standard growth at 30°C, Atg8-associated fluorescence was minimal, whereas fluorescence intensity increased markedly at 37°C (Fig. 3c). In contrast, incubation at 37°C plus 5% CO_2_ did not further enhance the signal, indicating that autophagosome-associated fluorescence responds primarily to elevated temperature rather than to CO_2_.

Golgi markers displayed a distinct but related response pattern. Both Cap6 and Ktr3 showed reduced fluorescence intensity at 37°C compared with 30°C; however, addition of 5% CO_2_ produced a modest increase in Golgi-associated signal relative to 37°C alone (Fig. 3c).

Although Golgi fluorescence did not return to 30°C levels, these data indicate that Golgi compartments are primarily temperature-sensitive, with CO_2_ exerting a partial modulatory effect.

In contrast to the responsive organelles described above, several markers showed no significant changes in fluorescence intensity across temperature and CO_2_ conditions. The mitochondrial marker Mjr1, the plasma membrane marker Pma1, and the vacuolar marker Vcx1 remained largely unchanged at 37°C and at 37°C plus 5% CO_2_ relative to 30°C (Fig. 3d). These findings indicate that not all organelles undergo quantitative remodeling under host-like conditions and underscore the specificity of the observed responses.

Collectively, these results show that elevated temperature and CO_2_ drive organelle-specific, stimulus-dependent remodeling patterns rather than a uniform increase or decrease in fluorescence across the cell.

### Dynamic changes in organelle organization under virulence factor-inducing conditions

We next asked whether the integration of organelle markers affects production of the two major *Cryptococcus* virulence factors, melanin and capsule, and whether organelle features are altered during virulence factor formation. Under melanin-inducing conditions on Niger seed medium, all marker strains melanized comparably, indicating that marker integration did not impair melanin production (Fig. 4a). Similarly, under capsule-inducing conditions in RPMI medium, capsule formation was comparable across marker strains, with no obvious differences in capsule thickness (Fig. 4b). Importantly, organelle-specific fluorescence signals remained clearly resolved during melanin and capsule induction without auto-fluorescence, demonstrating that marker visibility is maintained across distinct virulence factor–inducing environments (Fig. 4b).

**Fig. 4.**
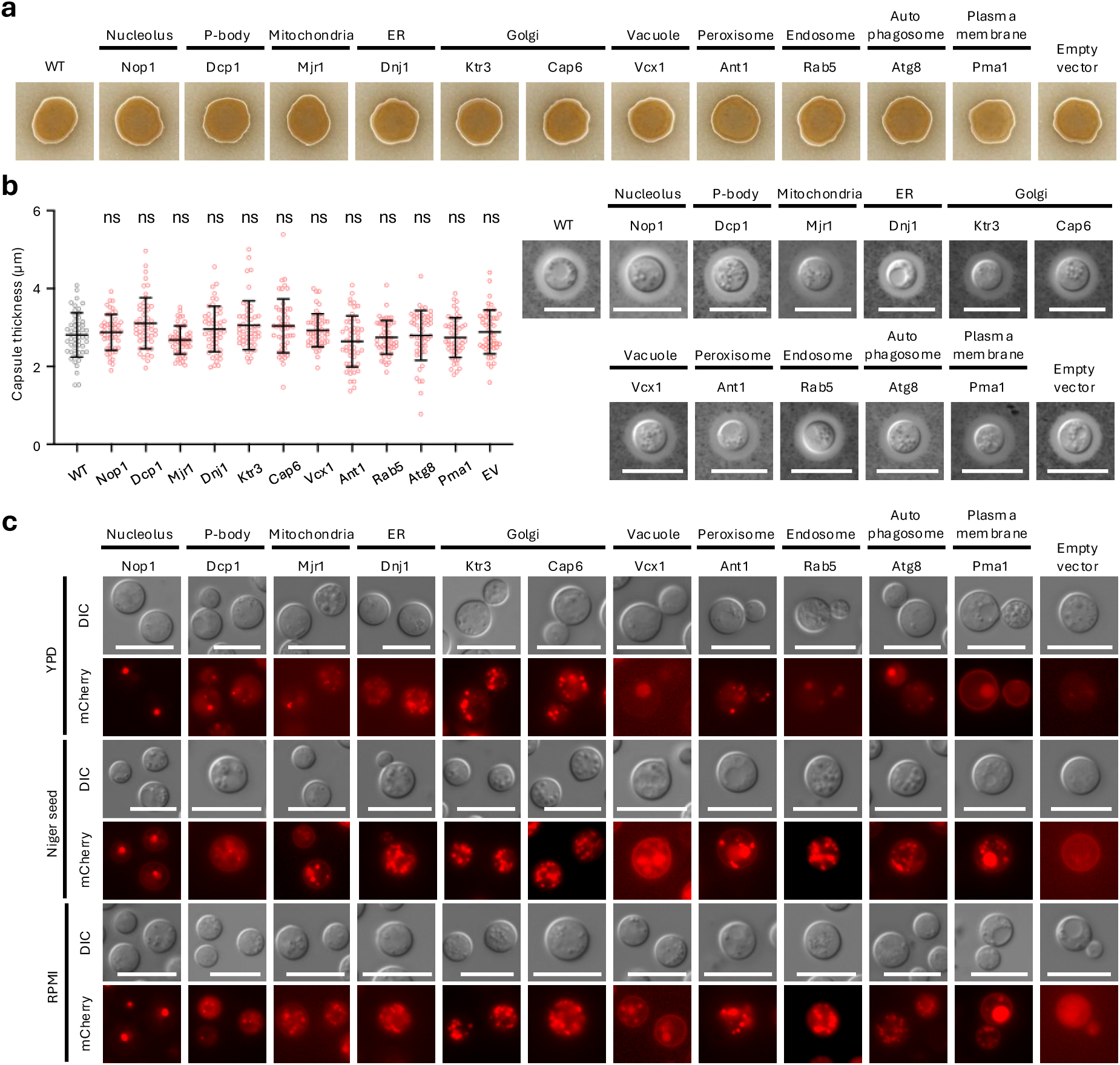
mCherry-tagged cellular organelle markers during virulence factor formation in *C. neoformans*. a and b) *C. neoformans* strains expressing mCherry-tagged organelle markers were initially cultured in YPD and subsequently spotted onto Niger seed agar or RPMI medium to induce melanin and capsule production, respectively. Plates were incubated at 37°C for 2 days and photographed to assess virulence factor formation. Capsule thickness was visualized and quantitatively measured. Capsule measurements were obtained from 50 individual cells per strain (*n* = 50). Data are presented as mean ± SEM. Statistical significance was evaluated using one-way ANOVA followed by Bonferroni’s multiple-comparison test (ns, not significant). Scale bars: 10 μm. c) For fluorescence analysis, cells were grown on YPD, Niger seed agar, and RPMI plates at 37°C for 2 days. Following incubation, cells were collected from the plates, resuspended in water, and subjected to fluorescence microscopy. Fluorescence images were captured using a fixed exposure time for each strain.

Notably, vesicle-associated compartments, including endosomes, autophagosomes, and vacuoles, were more prominent and exhibited clearer spatial organization under virulence factor–inducing conditions than during growth in YPD at 37°C, with compartment-specific localization patterns were maintained (Fig. 4c). In particular, endosomal and autophagosomal markers exhibited more clearly defined and spatially organized structures under these conditions, facilitating visualization of vesicle-associated compartmental architecture. The coordinated prominence of endosomal, autophagosomal, and vacuolar signals highlights vesicle trafficking–related compartments as defining features of cells under host-relevant, virulence-associated states (Fig. 4c).

Together, these observations show that the organelle marker set perform robustly across diverse host-relevant environments and identify vesicle-associated compartments as prominent features of *C. neoformans* during virulence factor induction.

### Distribution and inheritance of organelles during hyphal formation and sexual development

Sexual development in *Cryptococcus* proceeds through an ordered series of morphological transitions that are tightly coordinated with cellular differentiation. The process begins in the yeast-form and is initiated by the formation of conjugation tubes and then cell-cell fusion between compatible mating partners (**a** and α). These early events give rise to hyphae, during which dikaryotic hyphae elongate through polarized growth. Differentiation at hyphal tips can then produce basidia, specialized sexual structures in which karyogamy and meiosis occur. Finally, basidia generate chains of basidiospores, completing the sexual cycle and producing infectious propagules (Kronstad et al. 2011). This stepwise progression from yeast cells to hyphae that then produce basidia and basidiospores provides a structured framework to visualize dynamic cellular processes during sexual development.

To examine organelle dynamics during sexual development, genes representing individual organelles were GFP tagged and introduced into the *MAT***a** KN99 strain (Supplementary Figs. 2a and b). During zygote formation, *MAT*α cells typically extend conjugation tubes toward *MAT***a** cells, culminating in cell fusion (Gyawali and Lin 2013). After fusion, the *MAT*α nucleus is transferred into the *MAT***a** cell following cell fusion, where hyphal formation is initiated. Consistent with previous reports, the nucleolar markers Nop1–mCherry and Nop1–GFP converged within a single yeast cell (*MAT***a**) after fusion, indicating that both nuclei reside within the fused zygote (Fig. 5b). In contrast, other organelle markers displayed distinct inheritance behaviors. Markers for the ER, Golgi, peroxisomes, and the plasma membrane/vacuole system originating from *MAT***a** (GFP-tagged) cells were not confined to the *MAT***a** cell body but were detected within the conjugation tube and translocated into the opposing mating type cell (Fig. 5c-f). At this stage, endosomal markers did not appear as discrete vesicular structures; instead, both mCherry- and GFP-tagged endosomal signals were broadly distributed throughout the fused cytoplasm (Fig. 5g). Similarly, *MAT*α-derived GFP signals were detected in both parental cell bodies after fusion. Together, these observations indicate that following fusion of opposite mating type cells, organelles from both parents become extensively intermixed, reflecting not only plasma membrane continuity but also integration of intracellular compartments within the zygote.

**Fig. 5.**
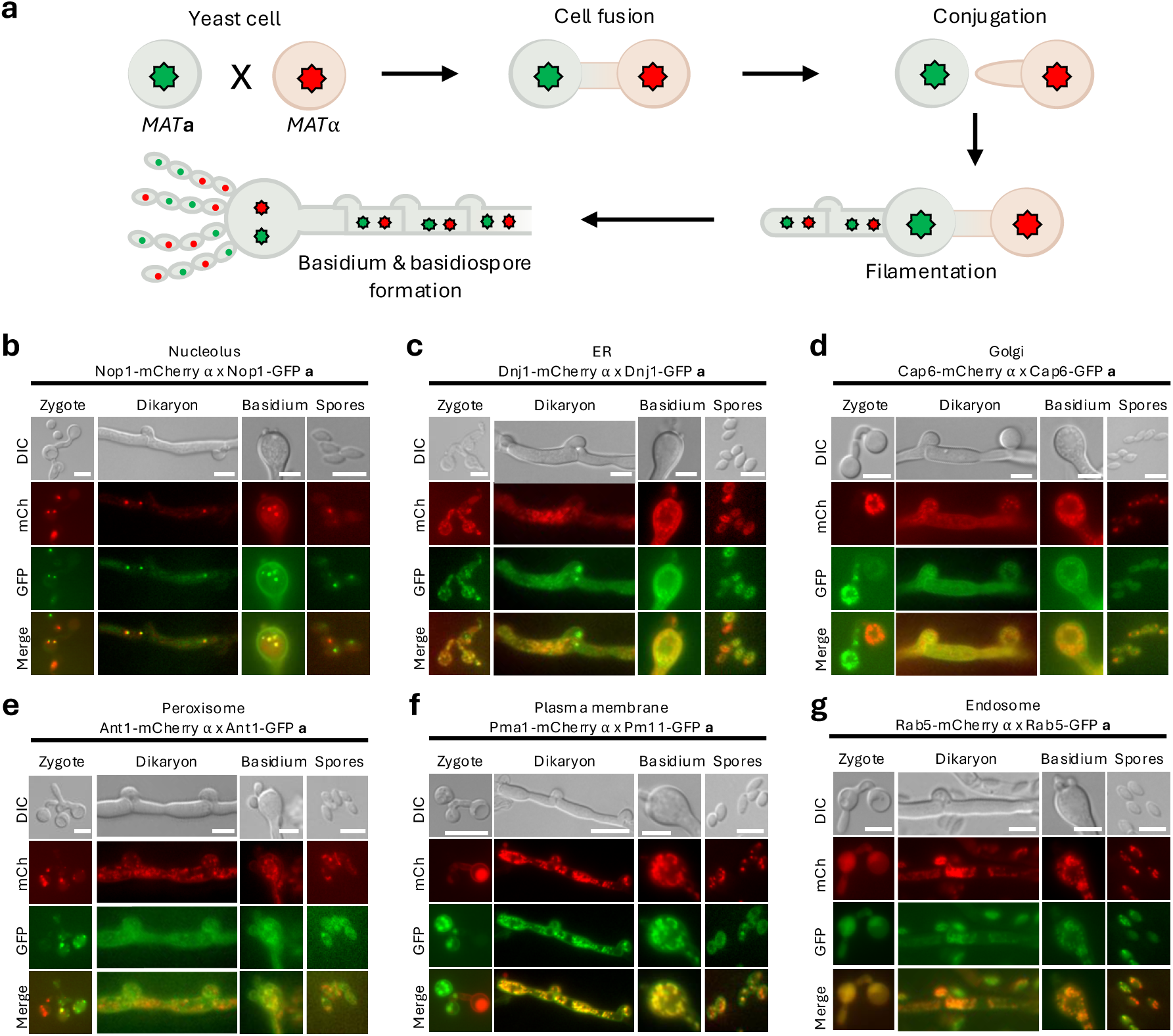
Organelle marker distribution during mating-induced filamentation in *C. neoformans*. a) The diagram illustrates the mating process between opposite mating types, including cell fusion, dikaryotic hyphal growth, clamp connection formation, basidium development, and basidiospore formation. b) Distribution and inheritance of intracellular organelles during mating visualized using fluorescent organelle markers. Strains of opposite mating types harboring identical organelle markers, each tagged with either mCherry or GFP to label the nucleolus, ER, Golgi, peroxisome, plasma membrane, and endosome, were crossed on MS medium to induce mating and filamentation. Fluorescence signals were visualized during hyphal, basidium, and basidiospore development, capturing both red and green organelle localization within filamentous cells. Representative panels include brightfield, individual fluorescence channels, and merged images. Scale bar; 5 µm

During dikaryotic hyphal growth, clamp connections formed normally, indicating that sexual development proceeded as expected. Both mCherry- and GFP-tagged organelle markers were broadly distributed throughout dikaryotic hyphae, consistent with extensive spreading of parental organelles after cell fusion (Fig. 5c-g). Signals from both markers were also detected in clamp cells, indicating that organelles are incorporated during clamp formation. For the ER marker, the GFP signal was occasionally observed in clamp cells without a corresponding mCherry signal (Fig. 5c). However, this pattern was not specific to the ER, as mCherry- and GFP-tagged ER signals were also frequently detected together within clamp connections. Similar to nuclei, which redistribute through septa and clamp connections during dikaryotic growth, organelles were likewise observed to pass through clamp connections into subsequent hyphal compartments. Golgi markers exhibited a distinct spatial organization, localizing outside the rounded, nucleus-like structures within clamp cells, consistent with perinuclear Golgi positioning. In addition, clamp connections displayed a polarized morphology reminiscent of yeast cells, further supporting coordinated cellular organization during dikaryotic growth (Fig. 5d).

As in dikaryotic hyphae, organelle markers from both mating partners were extensively interspersed throughout the developing basidia. Both mCherry- and GFP-tagged signals were broadly distributed across the basidial cytoplasm, indicating continued mixing of parental organelles during basidium development (Fig. 5b-g). For the nucleolar marker, progression through meiosis was readily observed within basidia (Fig. 5b). The ER marker prominently surrounded meiotic nuclei, forming well-defined circular structures that excluded the nuclear region (Fig. 5c). Golgi markers showed a similar organization, localizing to discrete basidial regions while remaining excluded from nuclear-associated areas (Fig. 5d). In contrast, peroxisomal and vacuolar markers, including the plasma membrane–associated vacuolar signal, did not show pronounced enrichment around the nuclei and instead appeared more broadly distributed within the basidium (Fig. 5e and f). Endosomal markers were detected as numerous small puncta throughout the basidial cytoplasm, consistent with vesicular organization. Notably, despite extensive mixing of parental organelles, basidia displayed clear spatial polarization of multiple organelle classes. Organelle markers were maintained throughout basidiospore formation and remained readily detectable in mature basidiospores, indicating efficient transmission of organelle labeling to sexual progeny (Fig. 5b-g). Endosomal markers were particularly well resolved in basidiospores, appearing as distinct puncta consistent with their vesicular nature (Fig. 5g).

Together, these findings indicate that sexual development in *C. neoformans* involves not only coordinated nuclear events but also dynamic, highly organized inheritance of intracellular organelles across developmental stages.

### A versatile platform for simultaneous organelle marker co-tagging to define subcellular localization of proteins of interest

To demonstrate the utility of the organelle marker platform for validating subcellular localization of proteins of interest, we used two complementary co-tagging strategies. These approaches enable visualization of candidate proteins relative to defined organelle markers either by generating diploids through mating or by introducing marker constructs directly into the same strain.

In the first approach, organelle marker strains of one mating type were crossed with strains of the opposite mating type expressing a GFP-tagged protein of interest to generate diploid cells, enabling simultaneous visualization of both signals. For this analysis, we selected Sua5, which has been reported to localize to mitochondria (Choi et al. 2024). *SUA5* was integrated at the safe haven locus with a backbone plasmid carrying the constitutive *H3* promoter and GFP. A *MAT*α strain harboring an mCherry-labeled mitochondrial marker and a *MAT***a** strain expressing Sua5-GFP were co-cultured on V8 medium for 24 h to induce mating, followed by selection on medium containing both markers (neomycin/G418 and hygromycin B) to isolate diploid cells.

As expected, Sua5-GFP colocalized with the mitochondrial marker, consistent with previous reports (Choi et al. 2024). In addition to mitochondrial localization, Sua5-GFP displayed a peripheral reticular pattern near mitochondria, suggesting association with an additional compartment (Fig. 6a). Sua5-GFP exhibited a perinuclear localization pattern, prompting examination of potential co-localization with ER and Golgi markers. To further define Sua5 localization, we generate diploids by crossing the Sua5-GFP strain with ER or Golgi marker strains. In these diploids, Sua5-GFP overlapped with the ER marker, whereas no comparable overlap was observed with Golgi markers (Fig. 6a). These results show that diploid-based colocalization provides a practical means to determine protein localization when constructs are in different mating types.

**Fig. 6.**
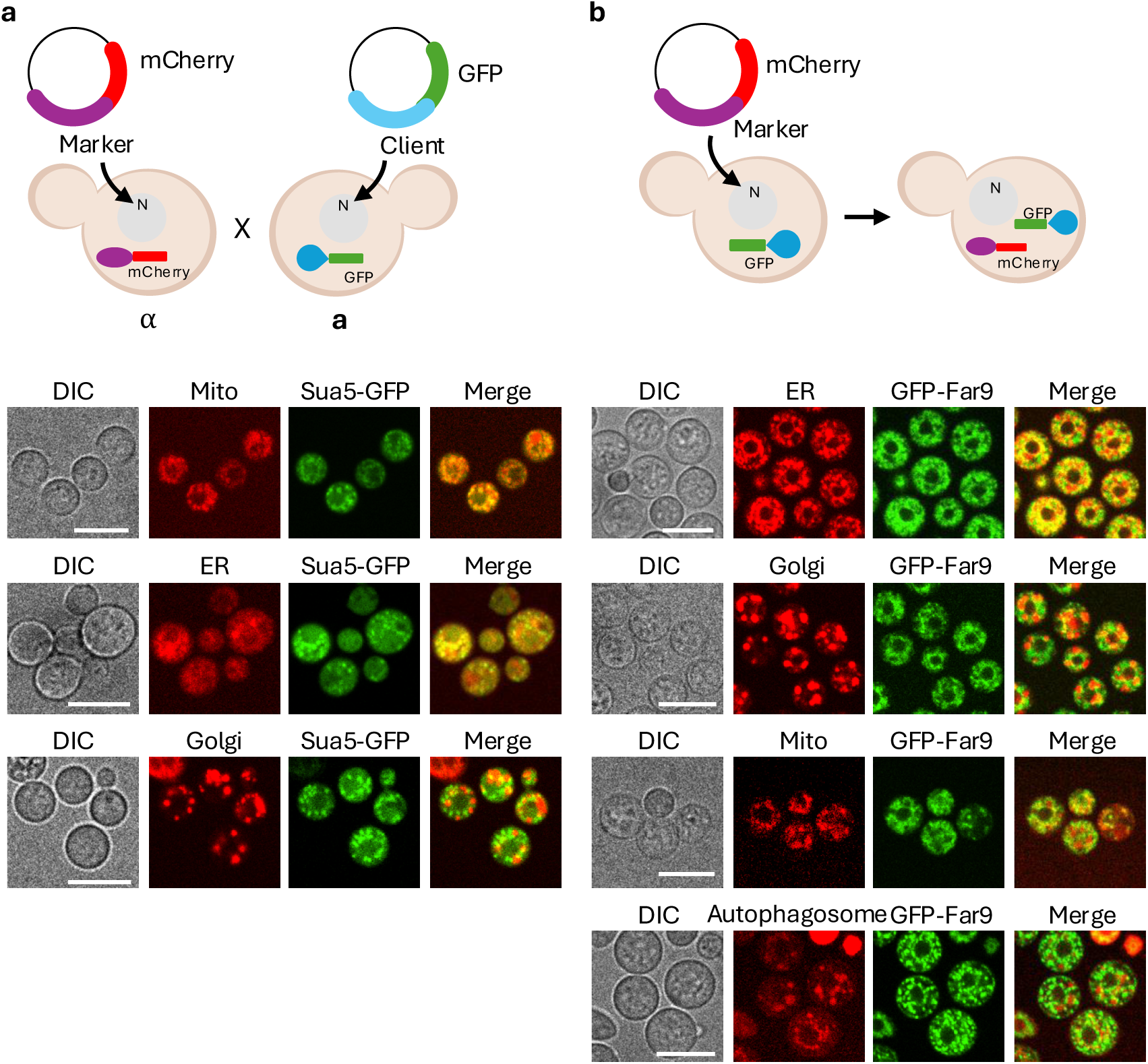
Co-localization of GFP-tagged proteins with mCherry-labeled organelle markers. a) Co-localization analysis in diploid cells generated by mating between opposite mating types. Opposite mating type strains were differentially labeled with mCherry or GFP and crossed to generate diploid cells. Upper schematics illustrate the experimental design, in which *MAT*α cells express mCherry-tagged organelle markers, while *MAT***a** cells express GFP-tagged client proteins, followed by cell fusion. Each mitochondrion (SYB11818), ER (YSB11832), and Golgi (YSC8) marker strains were crossed with the Sua5–GFP expressing strains (*MAT***a** SH:P*_H3_-GFP-HYG*, YSC106), and co-localization was analyzed in the resulting diploids. b) Co-tagging of two fluorescent markers in haploid yeast cells. Dual-color fluorescence labeling was achieved by introducing an mCherry-tagged organelle marker plasmid into strains expressing GFP-tagged proteins. Upper schematics depict the strategy in which mCherry-tagged organelle marker constructs were integrated into the safe haven locus of cells expressing GFP-tagged proteins. Cells expressing N-terminally GFP-tagged Far9 (*MAT*⍺ *far9*Δ::*NAT*, YSB11754) were individually transformed with plasmids encoding mCherry-tagged markers for the ER, Golgi, mitochondria, and autophagosomes, enabling co-localization analysis within a single haploid cell. For autophagosome visualization, cells were incubated at 37°C prior to imaging. Images were acquired using a confocal fluorescence microscope.

In the second approach, organelle marker plasmids were introduced directly into strains expressing a protein of interest to assess localization within a single genetic background. With this strategy, we examined Far9, a component of the STRIPAK complex reported to associate with membrane-linked compartments (Peterson et al. 2024; Peterson and Heitman 2025). We generated an N-terminally GFP-tagged Far9 construct and examined the localization of GFP-Far9 in *C. neoformans*. Organelle marker plasmids for the Golgi apparatus (Ktr3), ER (Dnj1), autophagosomes (Atg8), and mitochondria (Mjr1) were independently transformed into the *far9*Δ::*NAT GFP-FAR9-T_FAR9_-NEO* strain. Colocalization and spatial relationships between GFP-Far9 and each organelle marker were assessed by fluorescence microscopy. GFP-Far9 showed strong spatial association with the ER marker, with adjacent and partially overlapping localization. In contrast, GFP-Far9 did not show clear colocalization with mitochondrial, Golgi, or autophagosomal markers; instead, these signals were spatially adjacent or interspersed without consistent overlap (Fig. 6b), consistent with membrane tethering near, but not within, these compartments.

Together, these two co-tagging strategies demonstrate the versatility of the organelle marker platform for validating protein subcellular localization. Diploid-based co-tagging enables comparative localization analysis across mating types (and has the added advantage that diploid cells are larger than haploid cells, facilitating protein localization based on direct fluorescence microscopy), whereas direct co-introduction of markers allows assessment of spatial relationships within a single constructed strain, providing complementary approaches to resolve protein localization in *C. neoformans*.

## Discussion

In this study, we establish a comprehensive and versatile platform of fluorescent organelle markers for *C. neoformans* that enables visualization of subcellular organization across a broad range of physiological, developmental, and host-relevant conditions. By systematically validating marker localization under elevated temperature and CO_2_, in capsule-and melanin-inducing media, and throughout mating and basidial development, we show that organelle architecture in *Cryptococcus* is highly dynamic and strongly shaped by the environment. These findings underscore the value of examining subcellular organization beyond standard laboratory conditions, particularly in pathogenic fungi whose success depends on rapid adaptation to diverse host environments. Importantly, this platform also provides a flexible framework for visualizing proteins of interest alongside defined organelle markers, enabling co-localization analyses that support functional inference for uncharacterized proteins.

Compared with chemical dyes and organelle-specific trackers, genetically encoded fluorescent tagging offers several advantages for live-cell imaging and mechanistic studies. Protein-based tags enable visualization of organelles in living cells without fixation, eliminating the need for additional dye incubation and wash steps. Notably, dye-loading often requires incubation at elevated temperature, which can alter the properties of temperature-responsive proteins and confound phenotypic readouts. For example, whereas MitoTracker dyes label mitochondria in a membrane potential-dependent manner and therefore preferentially mark metabolically active mitochondria, Mjr1-based tagging uniformly labels the mitochondrial network, including less active regions and tubular structures. As a result, Mjr1 tagging allows robust assessment of mitochondrial organization and spatial distribution even under conditions in which mitochondrial activity is compromised. Together, these features highlight genetically encoded tagging as a less perturbative and more reliable approach than chemical dyes for interrogating intracellular organization and dynamics in live cells.

Previous work by Shi and Lin developed fluorescent tools to visualize subcellular compartments in *Cryptococcus* (Shi and Lin 2024). While that study established an important foundation for organelle imaging, our platform expands and complements those efforts in several ways. First, we incorporated additional markers for the vacuole, P-bodies, and autophagosomes, thereby broadening coverage of subcellular compartments implicated in stress adaptation, nutrient sensing, and intracellular recycling. Second, although both studies included endosomal markers, our work focuses on early endosomes, whereas their system visualized late endosomes, providing complementary resolution of endocytic trafficking. A further distinction is the marker design strategy. Shi and Lin utilized signal sequence–based targeting of fluorescent proteins, which can minimize perturbation of endogenous protein expression and may help preserve global proteostasis. In contrast, our approach tags well-characterized endogenous proteins that are representative of specific organelles. Although overexpression or tagging of native proteins can, in principle, influence cellular physiology or organelle dynamics, this strategy offers several advantages. By directly labeling bona fide organelle-resident proteins, our system increases confidence in organelle identity and enables reliable tracking of organelle morphology and remodeling. In addition, it facilitates interpretation of protein localization by providing established cellular landmarks for colocalization analyses. Thus, rather than relying solely on targeting sequences, our platform anchors localization studies within native subcellular architecture, providing a strong framework for defining protein distribution across organelles.

Utilizing this platform, we observed condition-dependent remodeling of multiple subcellular compartments across diverse environmental cues. Notably, increased nuclear and ER-associated signals coupled with reduced Golgi signal at elevated temperature suggests that transcriptional and translational output is not necessarily matched by downstream secretory maturation under stress. One explanation could be activation of ER quality control and unfolded protein response (UPR) pathways, which promote retention of newly synthesized proteins in the ER and limit ER-to-Golgi export during thermal stress (Cheon et al. 2014; Satpute-Krishnan et al. 2014). Alternatively, elevated temperature may induce structural and functional reorganization of the Golgi, such as fragmentation or reduced trafficking capacity, leading to an apparent decrease in Golgi-associated signal. Similar uncoupling between protein synthesis and secretory trafficking has been observed in other fungal systems adapting to environmental stress (Luo and Liu 2023). Together, these observations suggests that protein synthesis and secretory trafficking may be differentially coordinated in response to elevated temperature and CO_2_.

Under virulence factor-inducing conditions, vesicle trafficking–related compartments became more prominent, raising the question of why these pathways are preferentially engaged during pathogenic adaptation. One explanation is that vesicle trafficking is required for efficient secretion of major virulence determinants, including capsule components and extracellular enzymes, which depend on coordinated intracellular transport (de Oliveira et al. 2020; Rodrigues et al. 2007). In parallel, autophagy is often activated under host-like stresses and promotes survival by recycling cellular components and mitigating stress-induced damage (Jiang et al. 2020). Because many vesicle trafficking routes ultimately converge on the vacuole, a central hub for intracellular trafficking, the concomitant enhancement of vacuolar signals under these conditions is consistent with increased flux through endomembrane pathways. Together, these processes provide a plausible basis for the enhanced prominence of vesicle-associated compartments observed under virulence-associated conditions.

During sexual development, fusion between mating partners leads to mixing of cellular organelles. In contrast to the directional migration of nuclei, our observations suggest that organelles from both parents move and merge as the germ tube elongates. After cell fusion, organelles appeared to redistribute broadly during dikaryotic hyphal growth. Notably, organelles were also readily detected within clamp cells during septation and clamp connection formation, indicating that clamp cells are not organelle-poor or purely transient structures but instead contain established subcellular compartments.

Because clamp connections primarily ensure proper segregation of the two parental nuclei, the presence and organization of organelles within these structures raise questions about the relative timing of organelle versus nuclear inheritance. With the ER marker, we occasionally observed a single ER signal traversing the clamp connection into the adjacent compartment. This pattern suggests two non-mutually exclusive possibilities: nuclei may enter the clamp connection first and subsequently establish local organelle components through gene expression, or organelles may be delivered into the clamp connection in advance, analogous to organelle inheritance during vegetative budding, followed by nuclear migration. In the latter model, the detection of a single ER signal raises the possibility that ER derived from one mating type is preferentially transmitted at this stage.

In summary, we developed a versatile fluorescent organelle marker toolkit for *Cryptococcus*. This platform enables flexible analysis of protein subcellular localization with plasmid-based or strain-based approaches. Applying this system revealed condition-dependent changes in organelle organization across physiological, virulence-associated, and sexual development states, highlighting its utility for dissecting dynamic cellular processes in this pathogen.

## Data availability

Strains and plasmids described in this study can be provided upon request. All data supporting the conclusions of this study are presented within the manuscript, figures, and tables.

## Acknowledgments

We thank Dr. Patricia P. Peterson for sharing the *FAR9* sequence information, which facilitated this study. We also thank Dr. Jun Huang for comments on the manuscript and Dr. Claudia A. Petrucco for assistance with confocal microscopy.

## Author contributions

Y.C., Y.B., and J.H. conceived and designed the experiments, analyzed the data, and wrote and edited the manuscript. Y.C. performed the experiments.

---

**Figure S1.** Construction and functional validation of mCherry labeled organelle marker. a) Genotype validation of mCherry-tagged organelle marker strains. Correct integration of the mCherry tagging cassette at the safe haven locus was verified by diagnostic PCR, confirming both 5’-end and 3’-end homologous recombination events as illustrated in the schematic shown in Fig. 1b. All strains shown carry SH:P*_H3_*–gene–*mCherry* integrations, with individual genes encoding organelle markers fused to mCherry at the safe haven locus. The corresponding organelles and strain identifiers are indicated: *NOP1* (nucleolus, YSB11829), *DCP1* (P-body, YSB11825), *DNJ1* (ER, YSB11832), *KTR3* (Golgi, YSC8), *CAP6* (Golgi, YSC11), *VCX1* (vacuole, YSB11822), *RAB5* (endosome, YSC9), *MJR1* (mitochondria, YSB11818), *ANT1* (peroxisome, YSB11817), *ATG8* (autophagosome, YSB11835), and *PMA1* (plasma membrane, YSC4). “mCh” indicates the mCherry-only control strain (YSC1), while WT and DW represent the H99 wild-type strain and distilled water control, respectively. b) Gene expression analysis of genes flanking the safe haven locus and mCherry-tagged target genes. Transcript levels of genes located upstream and downstream of the safe haven integration site, as well as the corresponding mCherry-tagged target genes, were quantified by qRT-PCR. Expression levels were compared between the H99 wild-type strain (WT) and strains carrying SH:P*_H3_*-gene*-mCherry-HYG* integrations. Data are shown as relative expression normalized to WT. Bars represent the mean ± SEM. Statistical significance was assessed using one-way ANOVA with Bonferroni’s multiple-comparison test (ns, not significant; ***, *p* < 0.001; ****, *p* < 0.0001). c) Stress susceptibility assay. The susceptibility of the wild-type (WT) strain and mCherry-tagged cellular marker strains to antifungal drugs and other stress-inducing agents was examined. Cells were cultured overnight in YPD medium at 30°C, serially diluted (10-fold), and spotted onto YPD agar plates containing the indicated stressors. Abbreviations: HU, Hydroxy urea; MMS, methyl methanesulfonate; CR, Congo red; CFW, calcofluor white; TM, tunicamycin; DTT, dithiothreitol; tBOOH, tert-butyl hydroperoxide; H_2_O_2_, hydrogen peroxide; MD, menadione; FDX, fludioxonil; AmB, amphotericin B; Rapa, rapamycin.

**Figure S2.** Construction of GFP-tagged cellular marker strains. a) Genotype validation of GFP-tagged organelle marker strains. Correct integration of GFP-tagging cassettes at the safe-haven locus was confirmed by diagnostic PCR, verifying both 5’-and 3’-end homologous recombination events as illustrated in Fig. 1B. All strains carry SH:P*_H3_*-gene-GFP-*HYG* integrations, with individual genes encoding organelle markers fused to GFP at the safe haven locus (*NOP1*-GFP, YSC47; *DCP1-GFP*, YSC49; *DNJ1-GFP*, YSC52; *KTR3-GFP*, YSC136; *CAP6-GFP*, YSC178; *VCX1-GFP*, YSC179; *RAB5-GFP*, YSC159; *MJR1-GFP*, YSC44; *ANT1*-GFP, YSC156; *ATG8-GFP*, YSC162; and *PMA1*-*GFP*, YSC160). “GFP” denotes the GFP-only control strain, while WT and DW represent the KN99 wild-type strain and distilled water control, respectively. b) Representative fluorescence images of GFP-tagged organelle marker strains. All strains were cultured at 30°C and imaged live without fixation. For the strain expressing Dcp1-GFP, cells were grown at 30°C, shifted to 37°C for 15 min, and immediately imaged to induce P-body formation. For the autophagosome marker (Atg8) strain, cells were cultured in YNB medium supplemented with PMSF and nocodazole. Scale bars; 5 μm.

## Literature cited

1. Aboobakar, E.F., X. Wang, J. Heitman, and L. Kozubowski, 2011 The C2 domain protein Cts1 functions in the calcineurin signaling circuit during high-temperature stress responses in *Cryptococcus neoformans*. Eukaryot Cell 10 (12):1714–1723.

2. Aizer, A., Y. Brody, L.W. Ler, N. Sonenberg, R.H. Singer et al., 2008 The dynamics of mammalian P body transport, assembly, and disassembly in vivo. Mol Biol Cell 19 (10):4154–4166.

3. Altamirano, S., Z. Li, M.S. Fu, M. Ding, S.R. Fulton et al., 2021 The cyclin Cln1 controls polyploid titan cell formation following a stress-induced G(2) arrest in *Cryptococcus*. mBio 12 (5):e0250921.

4. Arras, S.D., J.L. Chitty, K.L. Blake, B.L. Schulz, and J.A. Fraser, 2015 A genomic safe haven for mutant complementation in *Cryptococcus neoformans*. PLoS One 10 (4):e0122916.

5. Bratton, E.W., N. El Husseini, C.A. Chastain, M.S. Lee, C. Poole et al., 2012 Comparison and temporal trends of three groups with cryptococcosis: HIV-infected, solid organ transplant, and HIV-negative/non-transplant. PLoS One 7 (8):e43582.

6. Casadevall, A., M.S. Fu, A.J. Guimaraes, and P. Albuquerque, 2019 The ’amoeboid predator-fungal animal virulence’ hypothesis. J Fungi (Basel) 5 (1).

7. Chadwick, B.J., L.C. Ristow, E.E. Blackburn, X. Xie, D.J. Krysan et al., 2025 Microevolution of *Cryptococcus neoformans* in high CO(2) converges on mutations isolated from patients with relapsed cryptococcosis. Cell Rep 44 (3):115349.

8. Cheon, S.A., K.W. Jung, Y.S. Bahn, and H.A. Kang, 2014 The unfolded protein response (UPR) pathway in *Cryptococcus*. Virulence 5 (2):341–350.

9. Choi, Y., H. Hyeon, K. Lee, and Y.S. Bahn, 2024 Sua5 catalyzing universal t(6)A tRNA modification is responsible for multifaceted functions of the KEOPS complex in *Cryptococcus neoformans*. mSphere 9 (1):e0055723.

10. Christopher, J.A., L.M. Breckels, O.M. Crook, M. Vazquez-Chantada, D. Barratt et al., 2025 Global proteomics indicates subcellular-specific anti-ferroptotic responses to ionizing radiation. Mol Cell Proteomics 24 (1):100888.

11. de Oliveira, H.C., R.F. Castelli, F.C.G. Reis, J. Rizzo, and M.L. Rodrigues, 2020 Pathogenic delivery: the biological roles of cryptococcal extracellular vesicles. Pathogens 9 (9).

12. Ding, H., M. Caza, Y. Dong, A.A. Arif, L.C. Horianopoulos et al., 2018 *ATG* genes influence the virulence of *Cryptococcus neoformans* through contributions beyond core autophagy functions. Infect Immun 86 (9).

13. El Magraoui, F A. Schrotter, R. Brinkmeier, L. Kunst, T. Mastalski et al., 2014 The cytosolic domain of Pex22p stimulates the Pex4p-dependent ubiquitination of the PTS1-receptor. PLoS One 9 (8):e105894.

14. Estrada, A.F., G. Muruganandam, C. Prescianotto-Baschong, and A. Spang, 2015 The ArfGAP2/3 Glo3 and ergosterol collaborate in transport of a subset of cargoes. Biol Open 4 (7):792–802.

15. Fan, Y., and X. Lin, 2018 Multiple applications of a transient CRISPR-Cas9 coupled with electroporation (TRACE) system in the *Cryptococcus neoformans* species complex. Genetics 208 (4):1357–1372.

16. Gyawali, R., and X. Lin, 2013 Prezygotic and postzygotic control of uniparental mitochondrial DNA inheritance in *Cryptococcus neoformans*. mBio 4 (2):e00112–00113.

17. Hammer, J.A., 3rd, and J.R. Sellers, 2011 Walking to work: roles for class V myosins as cargo transporters. Nat Rev Mol Cell Biol 13 (1):13–26.

18. Hein, M.Y., D. Peng, V. Todorova, F. McCarthy, K. Kim et al., 2025 Global organelle profiling reveals subcellular localization and remodeling at proteome scale. Cell 188 (4):1137–1155 e1120.

19. Horianopoulos, L.C., G. Hu, M. Caza, K. Schmitt, P. Overby et al., 2020 The novel J-domain protein Mrj1 is required for mitochondrial respiration and virulence in *Cryptococcus neoformans*. mBio 11 (3).

20. Horianopoulos, L.C., C.W.J. Lee, G. Hu, M. Caza, and J.W. Kronstad, 2021 Dnj1 promotes virulence in *Cryptococcus neoformans* by maintaining robust endoplasmic reticulum homeostasis under temperature stress. Front Microbiol 12:727039.

21. Hu, G., M. Caza, B. Cadieux, E. Bakkeren, E. Do et al., 2015 The endosomal sorting complex required for transport machinery influences haem uptake and capsule elaboration in *Cryptococcus neoformans*. Mol Microbiol 96 (5):973–992.

22. Jiang, S.T., A.N. Chang, L.T. Han, J.S. Guo, Y.H. Li et al., 2020 Autophagy regulates fungal virulence and sexual reproduction in *Cryptococcus neoformans*. Front Cell Dev Biol 8:374.

23. Keating, T.J., J.G. Peloquin, V.I. Rodionov, D. Momcilovic, and G.G. Borisy, 1997 Microtubule release from the centrosome. Proc Natl Acad Sci U S A 94 (10):5078–5083.

24. Kim, M.S., S.Y. Kim, J.K. Yoon, Y.W. Lee, and Y.S. Bahn, 2009 An efficient gene-disruption method in *Cryptococcus neoformans* by double-joint PCR with *NAT*-split markers. Biochem Biophys Res Commun 390 (3):983–988.

25. Kmetzsch, L., C.C. Staats, E. Simon, F.L. Fonseca, D.L. de Oliveira et al., 2010 The vacuolar Ca(2)(+) exchanger Vcx1 is involved in calcineurin-dependent Ca(2)(+) tolerance and virulence in *Cryptococcus neoformans*. Eukaryot Cell 9 (11):1798–1805.

26. Kozubowski, L., E.F. Aboobakar, M.E. Cardenas, and J. Heitman, 2011 Calcineurin colocalizes with P-bodies and stress granules during thermal stress in *Cryptococcus neoformans*. Eukaryot Cell 10 (11):1396–1402.

27. Kretschmer, M., J. Wang, and J.W. Kronstad, 2012 Peroxisomal and mitochondrial beta-oxidation pathways influence the virulence of the pathogenic fungus *Cryptococcus neoformans*. Eukaryot Cell 11 (8):1042–1054.

28. Kronstad, J.W., R. Attarian, B. Cadieux, J. Choi, C.A. D’Souza et al., 2011 Expanding fungal pathogenesis: *Cryptococcus* breaks out of the opportunistic box. Nat Rev Microbiol 9 (3):193–203.

29. Lee, S.C., and J. Heitman, 2012 Function of *Cryptococcus neoformans KAR7* (*SEC66*) in karyogamy during unisexual and opposite-sex mating. Eukaryot Cell 11 (6):783–794.

30. Li, S.C., and P.M. Kane, 2009 The yeast lysosome-like vacuole: endpoint and crossroads. Biochim Biophys Acta 1793 (4):650–663.

31. Lin, X., N. Chacko, L. Wang, and Y. Pavuluri, 2015 Generation of stable mutants and targeted gene deletion strains in *Cryptococcus neoformans* through electroporation. Med Mycol 53 (3):225–234.

32. Luo, A., and J.X. Liu, 2023 Rescuing the Golgi from heat damages by *ATG8*: restoration rather than clean-up. Stress Biol 3 (1):19.

33. Mansour, M.K., J.L. Reedy, J.M. Tam, and J.M. Vyas, 2014 Macrophage *Cryptococcus* interactions: an update. Curr Fungal Infect Rep 8 (1):109–115.

34. Mohamed, S.H., M.S. Fu, S. Hain, A. Alselami, E. Vanhoffelen et al., 2023 Microglia are not protective against cryptococcal meningitis. Nat Commun 14 (1):7202.

35. Mota, C., K. Kim, Y.J. Son, E.J. Thak, S.B. Lee et al., 2025 Evolutionary unique N-glycan-dependent protein quality control system plays pivotal roles in cellular fitness and extracellular vesicle transport in *Cryptococcus neoformans*. Elife 13.

36. Peterson, P.P., J.T. Choi, C. Fu, L.E. Cowen, S. Sun et al., 2024 The *Cryptococcus neoformans* STRIPAK complex controls genome stability, sexual development, and virulence. PLoS Pathog 20 (11):e1012735.

37. Peterson, P.P., and J. Heitman, 2025 The fungal STRIPAK complex: cellular conductor orchestrating growth and pathogenicity. PLoS Pathog 21 (9):e1013500.

38. Pope, R.E., and R.A. Prade, 2025 Vesicle-driven endomembrane systems in fungi. Microbiol Mol Biol Rev:e0029724.

39. Rodrigues, M.L., L. Nimrichter, D.L. Oliveira, S. Frases, K. Miranda et al., 2007 Vesicular polysaccharide export in *Cryptococcus neoformans* is a eukaryotic solution to the problem of fungal trans-cell wall transport. Eukaryot Cell 6 (1):48–59.

40. Satpute-Krishnan, P., M. Ajinkya, S. Bhat, E. Itakura, R.S. Hegde et al., 2014 ER stress-induced clearance of misfolded GPI-anchored proteins via the secretory pathway. Cell 158 (3):522–533.

41. Shi, R., and X. Lin, 2024 Illuminating the *Cryptococcus neoformans* species complex: unveiling intracellular structures with fluorescent-protein-based markers. Genetics 227 (3).

42. Sigaeva, A., C. Hutchings, A. Cesnik, K.S. Lilley, and E. Lundberg, 2026 Subcellular localization as a driver of protein function. Nat Rev Mol Cell Biol.

43. Stempinski, P.R., G.R. Gerbig, S.D. Greengo, and A. Casadevall, 2023 Last but not yeast-The many forms of *Cryptococcus neoformans*. PLoS Pathog 19 (1):e1011048.

44. Thak, E.J., Y.J. Son, D.J. Lee, H. Kim, J.H. Kim et al., 2022 Extension of O-linked mannosylation in the Golgi apparatus is critical for cell wall integrity signaling and interaction with host cells in *Cryptococcus neoformans* pathogenesis. mBio 13 (6):e0211222.

45. Wang, Y., P. Aisen, and A. Casadevall, 1995 *Cryptococcus neoformans* melanin and virulence: mechanism of action. Infect Immun 63 (8):3131–3136.

46. Xiao, W., H. Lu, B. Jiang, Y. Zheng, P. Chen et al., 2025 Virulence factors released by extracellular vesicles from *Cryptococcus neoformans*. Front Cell Infect Microbiol 15:1572520.

47. Zhao, X., W. Feng, X. Zhu, C. Li, X. Ma et al., 2019 Conserved autophagy pathway contributes to stress tolerance and virulence and differentially controls autophagic flux upon nutrient starvation in *Cryptococcus neoformans*. Front Microbiol 10:2690.

48. Zhao, Y.G., and H. Zhang, 2020 Phase separation in membrane biology: the interplay between membrane-bound organelles and membraneless condensates. Dev Cell 55 (1):30–44.

