## Supplementary Figs. 1-2 for "Fluorescent organelle markers in *Cryptococcus neoformans*: a versatile toolkit for live-cell subcellular localization"

**Figure S1 (Choi et al.)**

**a**

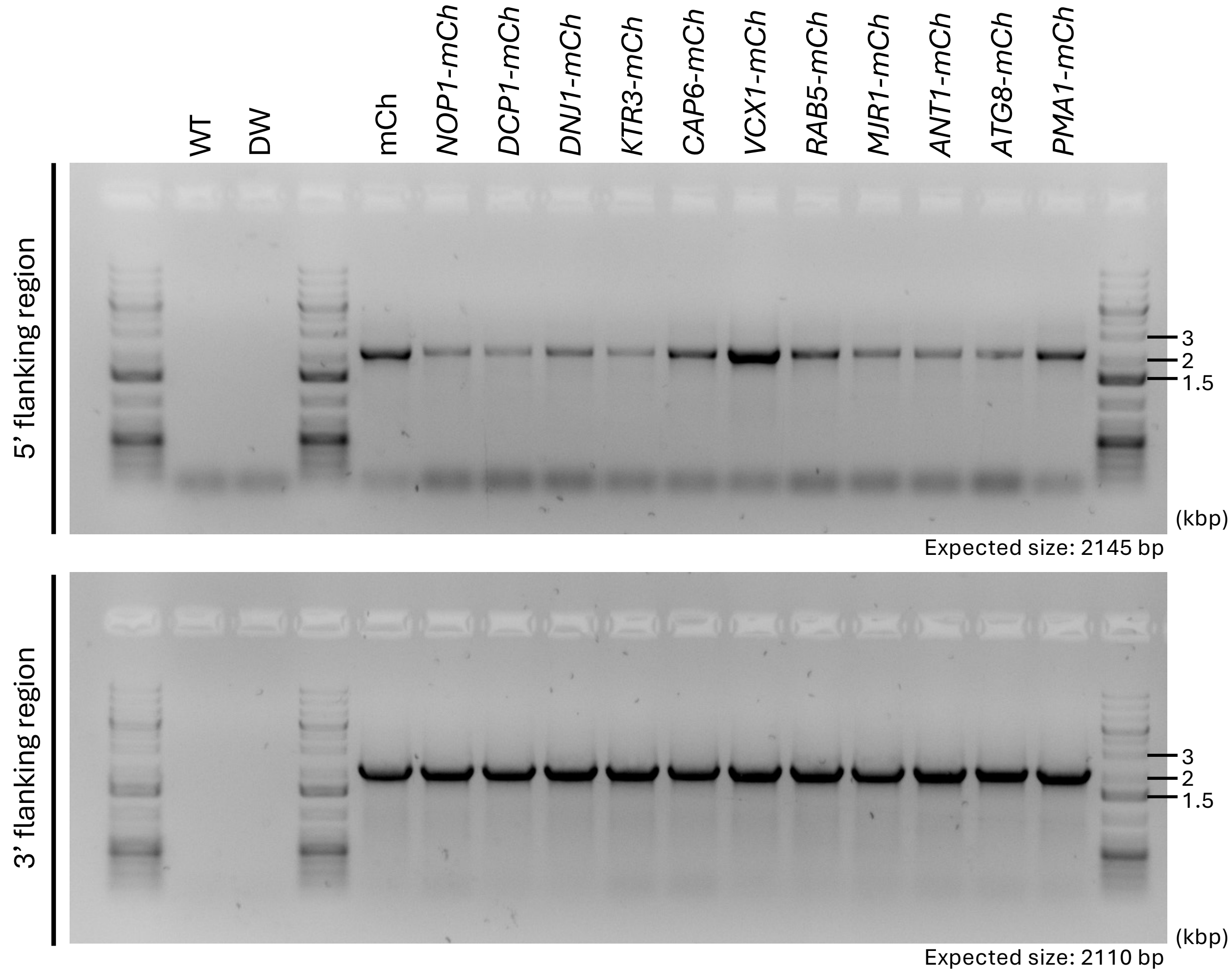


**Figure S1a. Genotype validation of mCherry-tagged organelle marker strains.**

### Figure S1 (Choi et al.)

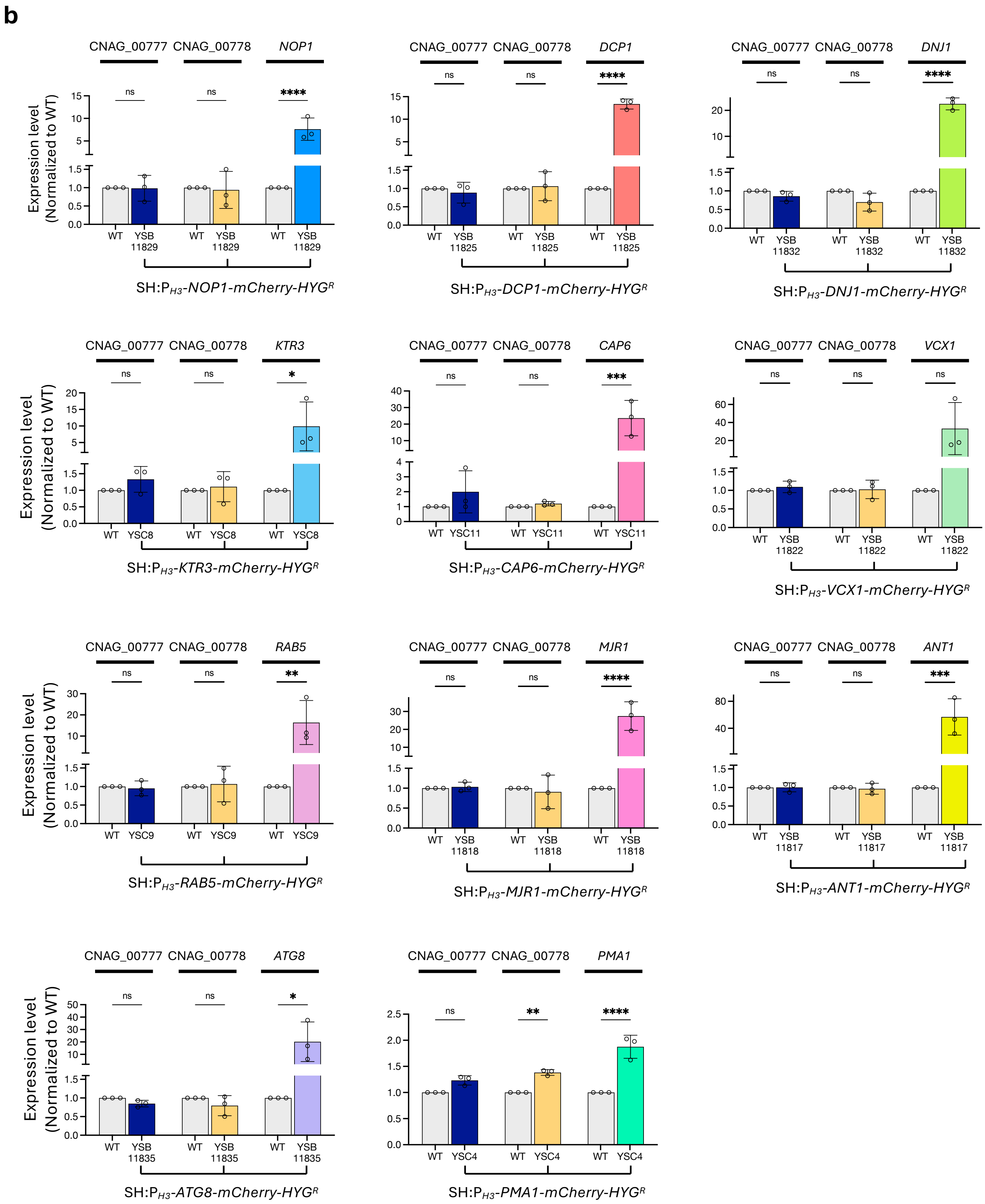


**Figure S1b. Gene expression analysis of genes flanking the safe haven locus and mCherry-tagged target genes.**

#### Figure S1 (Choi et al.)

c

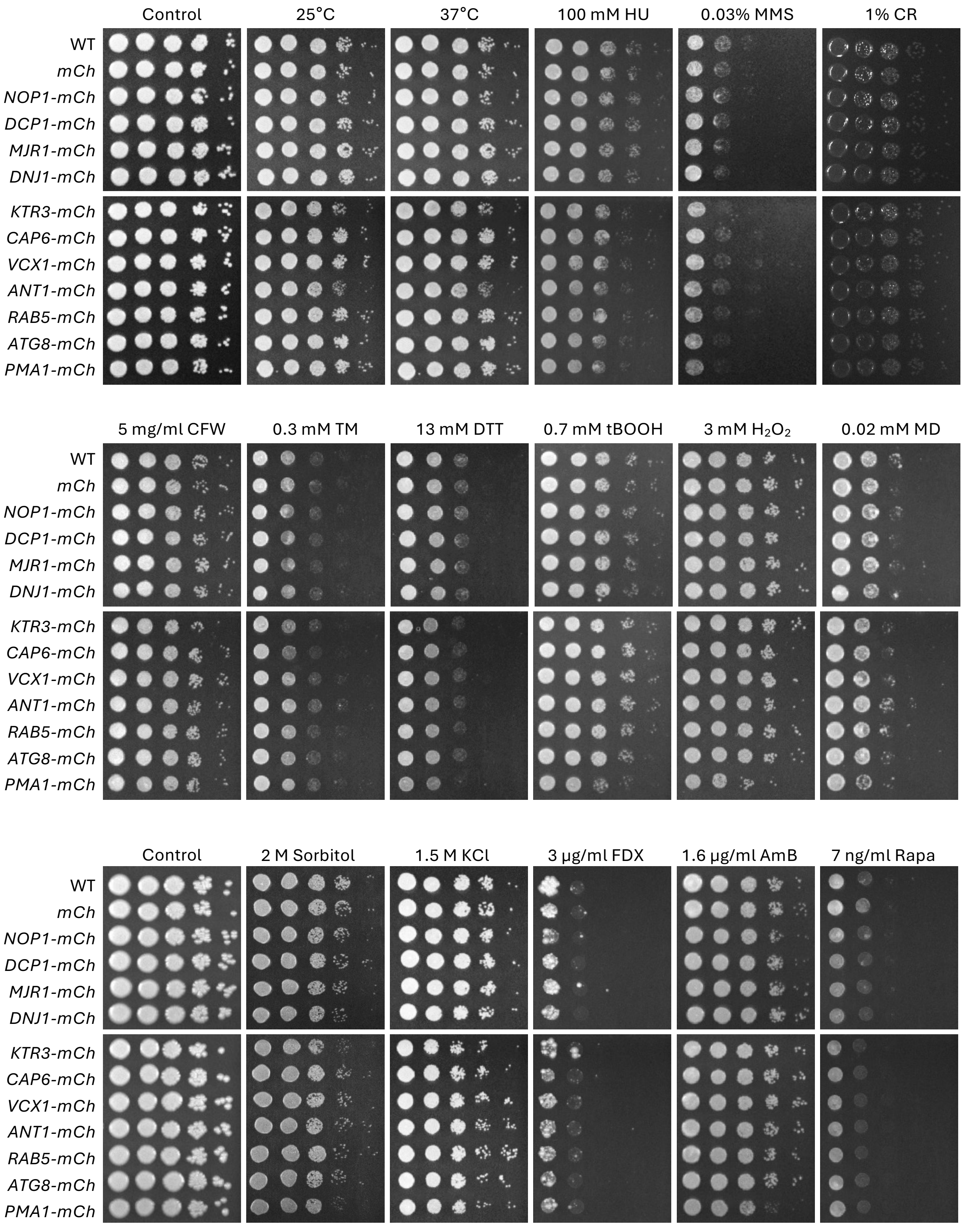


**Figure S1c. Stress susceptibility assay.**

#### Figure S2. (Choi et al.)

**a**

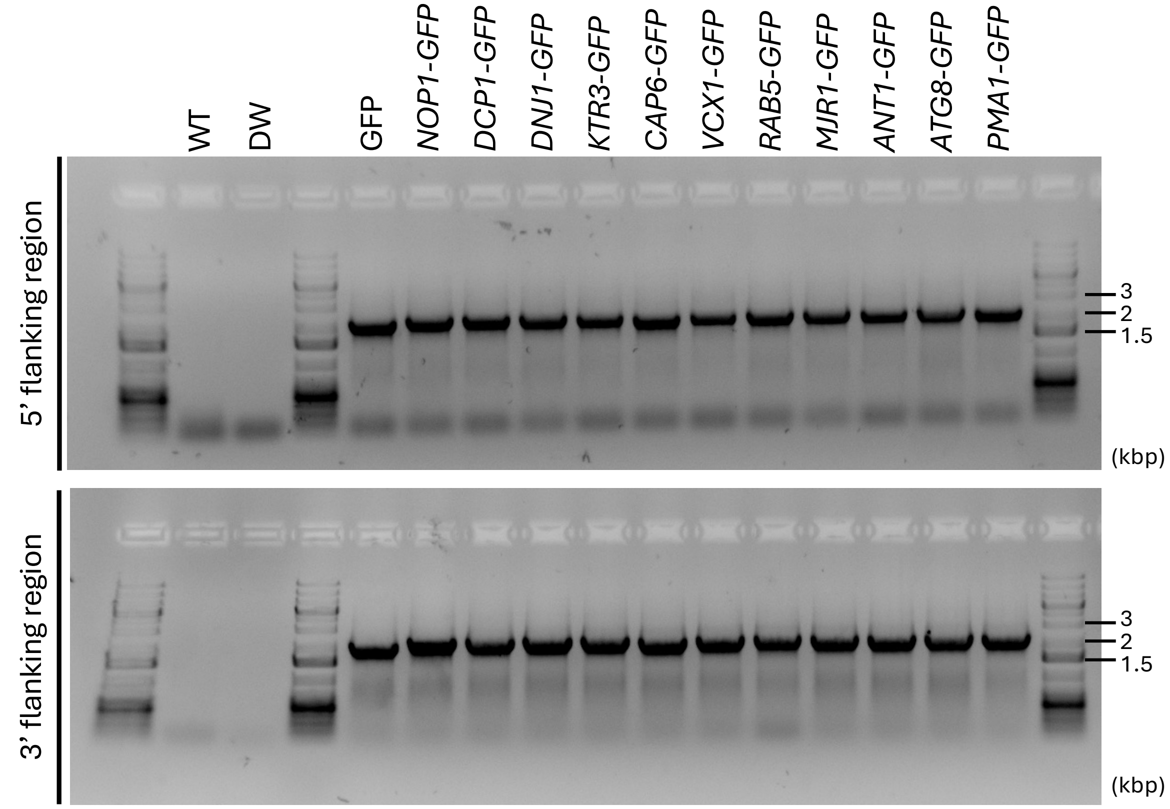


**b**

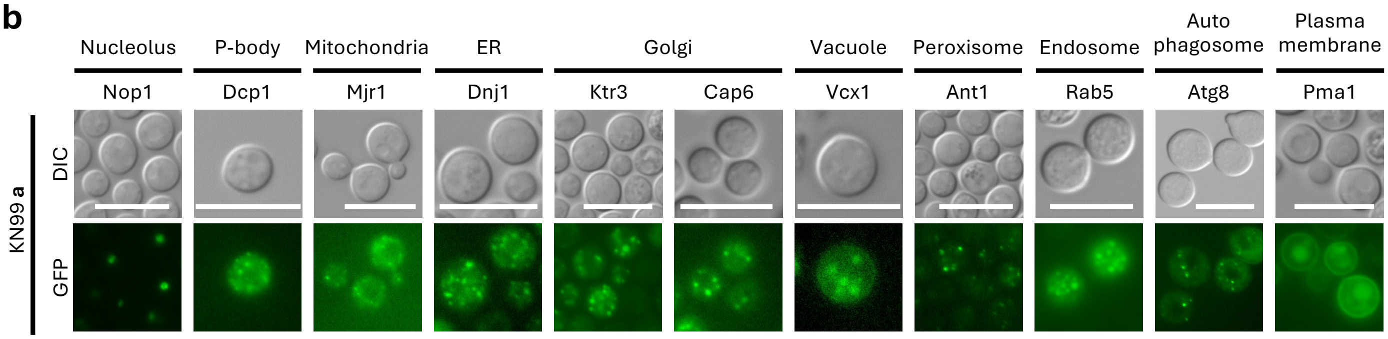


**Figure S2. Construction of GFP-tagged cellular marker strains.**
