## Supplementary material for "Fluorescent organelle markers in *Cryptococcus neoformans*: a versatile toolkit for live-cell subcellular localization": Table S1-3

**Table S1: Strains used in this study.**

| **Strain** | **Genotype** | **Parents** | **Reference** |
| --- | --- | --- | --- |
| H99 | *MATα* |  | (1) |
| KN99 | *MAT***a** |  | (2) |
| YSC1 | *MAT*α SH:P*_H3_-mCherry-HYG* | H99 | This study |
| YSB11829 | *MAT*α SH:P*_H3_-NOP1-mCherry-HYG* | H99 | This study |
| YSB11825 | *MAT*α SH:P*_H3_-DCP1-mCherry-HYG* | H99 | This study |
| YSB11832 | *MAT*α SH:P*_H3_-DNJ1-mCherry-HYG* | H99 | This study |
| YSC8 | *MAT*α SH:P*_H3_-KTR3-mCherry-HYG* | H99 | This study |
| YSC11 | *MAT*α SH:P*_H3_-CAP6-mCherry-HYG* | H99 | This study |
| YSB11822 | *MAT*α SH:P*_H3_-VCX1-mCherry-HYG* | H99 | This study |
| YSC9 | *MAT*α SH:P*_H3_-RAB5-mCherry-HYG* | H99 | This study |
| YSB11818 | *MAT*α SH:P*_H3_-MJR1-mCherry-HYG* | H99 | This study |
| YSB11817 | *MAT*α SH:P*_H3_-ANT1-mCherry-HYG* | H99 | This study |
| YSB11835 | *MAT*α SH:P*_H3_-ATG8-mCherry-HYG* | H99 | This study |
| YSC4 | *MAT*α SH:P*_H3_-PMA1-mCherry-HYG* | H99 | This study |
| YSC40 | MAT**a** SH:P*_H3_-GFP-HYG* | KN99 | This study |
| YSC47 | MAT**a** SH:P*_H3_-NOP1-GFP-HYG* | KN99 | This study |
| YSC49 | MAT**a** SH:P*_H3_-DCP1-GFP-HYG* | KN99 | This study |
| YSC52 | MAT**a** SH:P*_H3_-DNJ1-GFP-HYG* | KN99 | This study |
| YSC136 | MAT**a** SH:P*_H3_-KTR3-GFP-HYG* | KN99 | This study |
| YSC178 | MAT**a** SH:P*_H3_-CAP6-GFP-HYG* | KN99 | This study |
| YSC179 | MAT**a** SH:P*_H3_-VCX1-GFP-HYG* | KN99 | This study |
| YSC159 | MAT**a** SH:P*_H3_-RAB5-GFP-HYG* | KN99 | This study |
| YSC44 | MAT**a** SH:P*_H3_-MJR1-GFP-HYG* | KN99 | This study |
| YSC156 | MAT**a** SH:P*_H3_-ANT1-GFP-HYG* | KN99 | This study |
| YSC162 | MAT**a** SH:P*_H3_-ATG8-GFP-HYG* | KN99 | This study |
| YSC160 | MAT**a** SH:P*_H3_-PMA1-GFP-HYG* | KN99 | This study |
| YSC106 | MAT**a** SH:P*_H3_-SUA5-GFP-NEO* | KN99 | This study |
| YSC167 | *MAT*α SH:P*_H3_-DNJ1-mCherry-HYG* X *MAT****a*** SH:P*_H3_-SUA5-GFP-NEO* | YSB11832 x YSC106 | This study |
| YSC170 | *MAT*α SH:P*_H3_-MJR1-mCherry-HYG* X *MAT****a*** SH:P*_H3_-SUA5-GFP-NEO* | YSB11818 x YSC106 | This study |
| YSC176 | *MAT*α SH:P*_H3_-KTR3-mCherry-HYG* X *MAT****a*** SH:P*_H3_-SUA5-GFP-NEO* | YSC8 x YSC106 | This study |
| YSB11754 | *MAT*⍺ *far9*∆::*NAT* | H99 | (3) |
| YSC180 | *MAT*⍺ *far9*∆::*GFP-FAR9-*T*_FAR9_-NEO* | YSB11754 | This study |
| YSC185 | *MAT*⍺ *far9*∆::*GFP-FAR9-*T*_FAR9_-NEO* SH:P*_H3_*-*MJR1-mCherry-HYG* | YSC180 | This study |
| YSC186 | *MAT*⍺ *far9*∆::*GFP-FAR9-*T*_FAR9_-NEO* SH:P*_H3_*-*ATG8-mCherry-HYG* | YSC180 | This study |
| YSC187 | *MAT*⍺ *far9*∆::*GFP-FAR9-*T*_FAR9_-NEO* SH:P*_H3_*-*KTR3-mCherry-HYG* | YSC180 | This study |
| YSC194 | *MAT*⍺ *far9*∆::*GFP-FAR9-*T*_FAR9_-NEO* SH:P*_H3_*-*DNJ1-mCherry-HYG* | YSC180 | This study |

(1) Perfect, J.R., N. Ketabchi, G.M. Cox, C.W. Ingram, and C.L. Beiser, 1993 Karyotyping of *Cryptococcus neoformans* as an epidemiological tool. *J Clin Microbiol* 31 (12):3305–3309.

(2) Nielsen, K., G.M. Cox, P. Wang, D.L. Toffaletti, J.R. Perfect *et al*., 2003 Sexual cycle of *Cryptococcus neoformans* var. *grubii* and virulence of congenic a and alpha isolates. *Infect Immun* 71 (9):4831–4841.

(3) Peterson, P.P., J.T. Choi, C. Fu, L.E. Cowen, S. Sun *et al*., 2024 The *Cryptococcus neoformans* STRIPAK complex controls genome stability, sexual development, and virulence. *PLoS Pathog* 20 (11):e1012735.

**Table S2: Primers used in this study.**

| **Primer #** | **Sequence** | **Description** |
| --- | --- | --- |
| B21308 | ccacaacacatctatcacgc*ggccgc*ATGGCTTTCGGTGACAGAG | *NOP1* FWD |
| B21309 | acagagccaccgccacctgc*ggccgc*AGTGTGTCGTTGGTATATGC | *NOP1*_mCherry REV |
| JOHE54256 | atagagccaccgccacctgc*ggccgc*AGTGTGTCGTTGGTATATGC | *NOP1*_GFP REV |
| JOHE54268 | GTATGGCCAAGAAGCGAACC | *NOP1* qRT-LP |
| JOHE54269 | GCAAAGATGACGTCGACCAT | *NOP1* qRT-RP |
| JOHE54257 | ccacaacacatctatcacgc*ggccgc*ATGGCTTCAAGGTCCCCAAC | *DCP1* FWD |
| B21311 | acagagccaccgccacctgc*ggccgc*TCCCTCGTTCCACTGCCC | *DCP1*_mCherry REV |
| JOHE54258 | atagagccaccgccacctgc*ggccgc*TCCCTCGTTCCACTGCCC | *DCP1*_GFP REV |
| JOHE54270 | AGACGTCTCAAAGGTGCTCA | *DCP1* qRT-LP |
| JOHE54271 | ATGACAGCACGTGATTCTGC | *DCP1* qRT-RP |
| JOHE54259 | ccacaacacatctatcacgc*ggccgc*ATGGCGTTGGATAGAAGTGCC | *MJR1* FWD |
| B21313 | acagagccaccgccacctgc*ggccgc*CTCCCGGTGCGAAGGAGG | *MJR1*_mCherry REV |
| JOHE54260 | atagagccaccgccacctgc*ggccgc*CTCCCGGTGCGAAGGAGG | *MJR1-*GFP FWD |
| JOHE54272 | CAAGGATGGAGACCGTACGA | *MJR1* qRT-LP |
| JOHE54273 | CACCAACGACAGCAAGGAAT | *MJR1* qRT-RP |
| JOHE54051 | ccacaacacatctatcacgc*ggccgc*ATGAAAGGCTTTCTTCTAGTTG | *DNJ1* FWD |
| B21627 | acagagccaccgccacctgc*ggccgc*GTTCCACTGGAAGTGCATC | *DNJ1*-mCherry REV |
| JOHE54267 | atagagccaccgccacctgc*ggccgc*GTTCCACTGGAAGTGCATC | *DNJ1*-GFP REV |
| JOHE54282 | AGAACTGTCGCACCTCTTGA | *DNJ1* qRT-LP |
| JOHE54283 | CCAAAGCCTTAGTCGAGTGC | *DNJ1* qRT-RP |
| JOHE54020 | CCACAACACATCTATCACGCGGCCGCATGATGATTCCTCGCTGG | *KTR3* FWD |
| JOHE54021 | ACAGAGCCACCGCCACCTGCGGCCGCGAACATTTCTGTATATTTCTTAGTGC | *KTR3*-mCherry REV |
| JOHE54935 | atagagccaccgccacctgc*ggccgc*GAACATTTCTGTATATTTCTTAGTGC | *KTR3*-GFP REV |
| JOHE54660 | CTGGCTTCTTCTTCCGACAC | *KTR3* qRT_LP |
| JOHE54661 | TGACAAGGAAGGGGTCGTAG | *KTR3* qRT_RP |
| JOHE54022 | CCACAACACATCTATCACGCGGCCGCATGCCTCCCGACATACCTAAAC | *CAP6* FWD |
| JOHE54023 | ACAGAGCCACCGCCACCTGCGGCCGCAGATTGAACTGTAGTCTCAGAAGC | *CAP6*-mCherry REV |
| JOHE54934 | atagagccaccgccacctgc*ggccgc*AGATTGAACTGTAGTCTCAGAAGC | *CAP6*-GFP REV |
| JOHE54662 | TCAGGTTGCTGGTATGCTTG | *CAP6* qRT_LP |
| JOHE54663 | GCAGAGTCACCCTCGAAAAG | *CAP6* qRT_RP |
| JOHE54261 | ccacaacacatctatcacgc*ggccgc*ATGTTCCCCCAAAGCAAGTCC | *VCX1* FWD |
| B21475 | acagagccaccgccacctgc*ggccgc*GCCGGTGACGCTGGATGA | *VCX1*-mCherry REV |
| JOHE54262 | atagagccaccgccacctgc*ggccgc*GCCGGTGACGCTGGATGA | *VCX1*-GFP REV |
| JOHE54274 | GGGGAAGTATGAAAGCTGCG | *VCX1* qRT_LP |
| JOHE54275 | AGATGCTGTCCTTTCCACCA | *VCX1* qRT_RP |
| JOHE54263 | ccacaacacatctatcacgc*ggccgc*ATGGCACCTGTACACCATC | *ANT1 FWD* |
| B21621 | acagagccaccgccacctgc*ggccgc*TGTTTTGGCCAAGACGCG | *ANT1-mCherry REV* |
| JOHE54264 | atagagccaccgccacctgc*ggccgc*TGTTTTGGCCAAGACGCG | *ANT1-GFP REV* |
| JOHE54276 | AAGCAAAGGTCGTCGAATCG | *ANT1* qRT_LP |
| JOHE54277 | CATTCGCTAGGAGAGGCTGA | *ANT1* qRT_RP |
| JOHE54018 | CCACAACACATCTATCACGCGGCCGCATGTCTCGAACAACCTCG | *RAB5 FWD* |
| JOHE54019 | ACAGAGCCACCGCCACCTGCGGCCGCACATGTACAAGCTGAAAGAG | *RAB5-mCherry REV* |
| JOHE54936 | atagagccaccgccacctgc*ggccgc*ACATGTACAAGCTGAAAGAG | *RAB5-GFP REV* |
| JOHE54284 | GCGCTGCATTCCTTACTCAA | *RAB5* qRT_LP |
| JOHE54285 | TGGGAGCCAGTGACTTGTAG | *RAB5* qRT_RP |
| JOHE54265 | ccacaacacatctatcacgc*ggccgc*ATGGTCCGAAGCAAGTTTAAG | *ATG8 FWD* |
| B21623 | acagagccaccgccacctgc*ggccgc*CTCGGAGATGGCGTATTG | *ATG8-mCherry REV* |
| JOHE54266 | atagagccaccgccacctgc*ggccgc*CTCGGAGATGGCGTATTG | *ATG8-GFP REV* |
| JOHE54278 | CCAACAGCGGCACTTATGAG | *ATG8* qRT_LP |
| JOHE54279 | TTGTTCAAGGTCGCCAAAGG | *ATG8* qRT_RP |
| JOHE54932 | ccacaacacatctatcacgc*ggccgc*ATGTCTGACAACGAAAAAGTCGGTCACACCGAG | *PMA1 FWD* |
| B21625 | acagagccaccgccacctgc*ggccgc*CGCCGCGGGCCTGGAGTG | *PMA1-mCherry REV* |
| JOHE54933 | atagagccaccgccacctgc*ggccgc*CGCCGCGGGCCTGGAGTG | *PMA1-GFP REV* |
| JOHE54280 | TGGTCCTGATCATCGCTGTT | *PMA1* qRT_LP |
| JOHE54281 | TTCGGCAAGATCCCAAGAGT | *PMA1* qRT_RP |
| B21401 | GTGCAGATGGTCGCTCAGTA | Safe heaven upstream region diagnostic PCR SO1_LP |
| B21402 | CGTCAAACAAACATCCATGC | Safe heaven upstream region diagnostic PCR SO1_RP |
| B21403 | GGCCTCTTCGCTATTACGC | Safe heaven downstream region diagnostic PCR SO2_LP |
| B21404 | TTTTGAGCATTGACGCAACT | Safe heaven downstream region diagnostic PCR SO2_RP |
| JOHE54716 | ccctctagatgcatgctcgaACACGATGTTTTCCATCAG | SH-P_H3_ part FWD (for SH-H3-GFP-TER) |
| JOHE54150 | CTCCTCGCCCTTGCTCACCATAGAGCCACCGCCACCTGCGG | SH-P_H3_ part REV (for SH-H3-GFP-TER) |
| JOHE54151 | CCGCAGGTGGCGGTGGCTCTATGGTGAGCAAGGGCGAGGAG | GFP FWD (for SH-H3-GFP-TER) |
| JOHE54152 | GAAAGAAGAACGATTCTTTTAgTACAGCTCGTCCATGCCGTG | GFP REV (for SH-H3-GFP-TER) |
| JOHE54153 | CACGGCATGGACGAGCTGTAcTAAAAGAATCGTTCTTCTTTC | TER FWD (for SH-H3-GFP-TER) |
| JOHE54717 | tgcagatatccatcacactgGTACCAATCTATCCCTCTC | TER REV (for SH-H3-GFP-TER) |
| JOHE54286 | GTTCGAGACTTTCAATGCCC | ACT1_qLP |
| JOHE54287 | ACCAGAGTCAAGAACGATAC | ACT1 qRP |
| JOHE55518 | tagtaacggccgccagtgtg*ctgg*GCTGCGAGGATGTGAGCTG | NEO marker insertion FWD |
| JOHE55519 | gatagattggtaccagtgtg*atgg*GGTTTATCTGTATTAACACGGAAGAGATG | NEO marker insertion REV |
| JOHE55962 | ccacaacacatctatcacgc*ggccgc*ATGCGGGCAACTGTTCGC | pNEO_SH_PH3_SUA5_GFP FWD |
| JOHE55963 | atagagccaccgccacctgc*ggccgc*CGTAGAACTAACGTCCACCC | pNEO_SH_PH3_SUA5_GFP REV |
| JOHE55960 | gcgaagaggcccgcaccgat*cg*AGCCAAAGCGTCAGTTATATC | pNEO_5'_GFP_FAR9 5' FWD |
| JOHE57050 | gtgaacagctcctcgcccttgctcaccatGTTGAGTCGTTCGTAGTCAATGGTCTTCTC | pNEO_5'_GFP_FAR9 5' REV |
| JOHE57051 | GAGAAGACCATTGACTACGAACGACTCAACatggtgagcaagggcgaggagctgttcac | pNEO_5'_GFP_FAR9 GFP-FAR9-TER FWD |
| JOHE55961 | caactgttgggaagggcgat*cg*GATGCCAGCTCTGGGTAAG | pNEO_5'_GFP_FAR9 GFP-FAR9-TER REV |
| JOHE54848 | caccggcagggtatactgttGGACGTTTATCTTCTTGCTTgttttagagctagaaatagc | Safe heaven region gRNA |
| JOHE54910 | gtaaaacgacggccagt | M13F for CRISPR plasmid |
| JOHE54911 | caggaaacagctatgac | M13R for CRISPR plasmid |

**Table S3: Plasmids used in this study.**

| **Plasmid** | **Description** | **Bacterial Marker** | **Reference** |
| --- | --- | --- | --- |
| YSBE1510 | pHYG_SH_mCherry | Ampicillin | This study |
| YSBE1511 | pHYG_SH_P_H3_-mCherry | Ampicillin | This study |
| YSBE1512 | pHYG_SH_P_H3_-NOP1-mCherry | Ampicillin | This study |
| YSBE1513 | pHYG_SH_P_H3_-DCP1-mCherry | Ampicillin | This study |
| YSBE1514 | pHYG_SH_P_H3_-MJR1-mCherry | Ampicillin | This study |
| YSBE1515 | pHYG_SH_P_H3_-VCX1-mCherry | Ampicillin | This study |
| YSBE1516 | pHYG_SH_P_H3_-ANT1-mCherry | Ampicillin | This study |
| YSBE1517 | pHYG_SH_P_H3_-ATG8-mCherry | Ampicillin | This study |
| YSBE1518 | pHYG_SH_P_H3_-PMA1-mCherry | Ampicillin | This study |
| YSBE1519 | pHYG_SH_P_H3_-DNJ1-mCherry | Ampicillin | This study |
| YSCE1 | pHYG_SH_P_H3_-RAB5-mCherry | Ampicillin | This study |
| YSCE2 | pHYG_SH_P_H3_-CAP6-mCherry | Ampicillin | This study |
| YSCE3 | pHYG_SH_P_H3_-KTR3-mCherry | Ampicillin | This study |
| YSCE4 | pHYG | Ampicillin | This study |
| YSCE5 | pHYG_SH_P_H3_-GFP | Ampicillin | This study |
| YSCE6 | pHYG_SH_P_H3_-ANT1-GFP | Ampicillin | This study |
| YSCE8 | pHYG_SH_P_H3_-NOP1-GFP | Ampicillin | This study |
| YSCE9 | pHYG_SH_P_H3_-VCX1-GFP | Ampicillin | This study |
| YSCE10 | pHYG_SH_P_H3_-ATG8-GFP | Ampicillin | This study |
| YSCE11 | pHYG_SH_P_H3_-DNJ1-GFP | Ampicillin | This study |
| YSCE12 | pHYG_SH_P_H3_-MJR1-GFP | Ampicillin | This study |
| YSCE13 | pHYG_SH_P_H3_-DCP1-GFP | Ampicillin | This study |
| YSCE14 | pHYG_SH_P_H3_-RAB5-GFP | Ampicillin | This study |
| YSCE15 | pHYG_SH_P_H3_-CAP6-GFP | Ampicillin | This study |
| YSCE16 | pHYG_SH_P_H3_-KTR3-GFP | Ampicillin | This study |
| YSCE17 | pHYG_SH_P_H3_-PMA1-mCherry | Ampicillin | This study |
| YSCE18 | pNEO_SH_P_H3_-GFP | Ampicillin | This study |
| YSCE19 | pNEO_SH_P_H3_-SUA5-GFP | Ampicillin | This study |
| YSCE25 | pNEO_5'_FAR9__GFP_FAR9_T_FAR9_ | Kanamycin | This study |
| pBHM2329 | pRS316-P_CnU6_-scaffold | Ampicillin | (1) |
| pBHM2403 | pRS316_P_TEF1__CnoCas9 | Ampicillin | (1) |

(1) Huang, M.Y., M.B. Joshi, M.J. Boucher, S. Lee, L.C. Loza *et al*., 2022 Short homology-directed repair using optimized Cas9 in the pathogen *Cryptococcus neoformans* enables rapid gene deletion and tagging. *Genetics* 220 (1).
